# Insertion and Anchoring of HIV-1 Fusion Peptide into Complex Membrane Mimicking Human T-cell

**DOI:** 10.1101/2024.08.02.606381

**Authors:** Mingfei Zhao, Laura Joana Silva Lopes, Harshita Sahni, Anju Yadav, Hung N Do, Tyler Reddy, Cesar A. López, Chris Neale, S Gnanakaran

**Affiliations:** T-6 Theoretical Biology and Biophysics, Los Alamos National Laboratory, Los Alamos NM USA; Department of Computer Science, University of New Mexico, Albuquerque NM, USA; Department of Chemistry and Biochemistry, University of Texas at El Paso, El Paso TX, USA; CCS-7 Applied Computer Science Group, Los Alamos National Laboratory, Los Alamos NM USA

**Keywords:** HIV-1, fusion peptide, complex membrane, viral cell entry, multiscale modeling, molecular dynamics simulation, coarse-grained simulations

## Abstract

A fundamental understanding of how HIV-1 envelope (Env) protein facilitates fusion is still lacking. The HIV-1 fusion peptide, consisting of 15 to 22 residues, is the N-terminus of the gp41 subunit of the Env protein. Further, this peptide, a promising vaccine candidate, initiates viral entry into target cells by inserting and anchoring into human immune cells. The influence of membrane lipid reorganization and the conformational changes of the fusion peptide during the membrane insertion and anchoring processes, which can significantly affect HIV-1 cell entry, remains largely unexplored due to the limitations of experimental measurements. In this work, we investigate the insertion of the fusion peptide into an immune cell membrane mimic through multiscale molecular dynamics simulations. We mimic the native T-cell by constructing a 9-lipid asymmetric membrane, along with geometrical restraints accounting for insertion in the context of gp41. To account for the slow timescale of lipid mixing while enabling conformational changes, we implement a protocol to go back and forth between atomistic and coarse-grained simulations. Our study provides a molecular understanding of the interactions between the HIV-1 fusion peptide and the T-cell membrane, highlighting the importance of conformational flexibility of fusion peptides and local lipid reorganization in stabilizing the anchoring of gp41 into the targeted host membrane during the early events of HIV-1 cell entry. Importantly, we identify a motif within the fusion peptide critical for fusion that can be further manipulated in future immunological studies.

Table of Content.

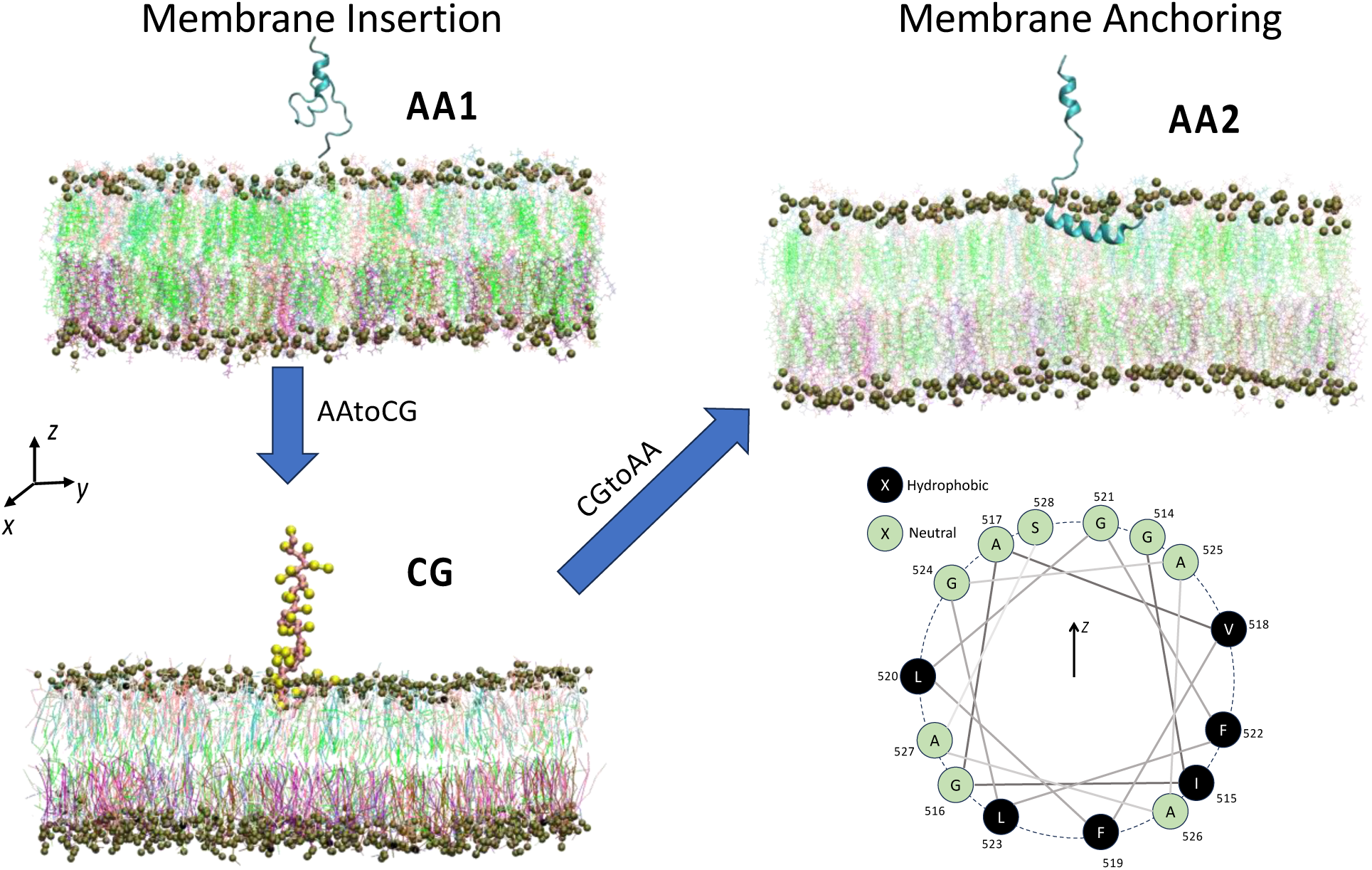

## 1. Introduction

The HIV-1 fusion peptide is the N-terminus of the gp41 subunit for the HIV-1 envelope protein (Env) and is responsible for initiating viral/host membrane fusion by inserting and anchoring it onto the host cell membrane.[1] As a result, HIV-1 fusion peptide, host cell membranes, and the interactions between HIV-1 fusion peptides and cell membranes draw significant attention from scientists for vaccine development and drug discovery to inhibit HIV-1 cell entry.[2–12] Moreover, the HIV-1 fusion peptide has been widely recognized as a promising vulnerable site to bind neutralizing antibodies.[5, 13, 14]

HIV-1 Env proteins undergo a series of conformational changes before inserting fusion peptide into cell membranes, transitioning from pre-fusion conformation to a partially open conformation after binding to a CD4 receptor and finally to the hypothesized pre-hairpin conformation.[15] In the pre-hairpin conformation, the gp41 heptad repeat 1 region coil is formed and exposed during full CD4-bound conformation.[15] As a result, the hydrophobic fusion peptide is exposed and inserted into the target host membrane, with gp41 forming a stable six-helical bundle to drive membrane fusion.[1, 15] The sequence of HIV fusion peptides is highly conserved among different variants of HIV-1 and is also similar to that of HIV-2 and SIV.[16, 17] This pre-hairpin conformation is critical for studying the membrane insertion of fusion peptides, but the detailed atom-level structure has not been revealed in experiments due to the rapid fusion process.[18–20] Recently, Ladinsky et al. presented the first direct visualization of membrane attachment sites with spokes representing the pre-hairpin intermediate through electron tomography.[18] Additionally, Lin and Da investigated the gp41 refolding dynamics from pre-fusion to pre-hairpin states by molecular dynamics simulation.[19] These experimental and computational studies agree that the fully extended gp41 helical bundle is the likely conformation before the fusion peptide inserts into the membrane.

Lipid compositions and lipid domains of target host membranes also play a role in the HIV-1 membrane fusion process. Tamm et al. have found that HIV-1 fusion peptides prefer to bind at the interface of liquid-ordered and liquid-disordered domains with different cholesterol concentrations due to line tension as a driving force.[21–23] Cholesterol can promote HIV-1 membrane fusion by modulating membrane properties,[24–26] conformation of fusion peptides,[4, 7] and membrane-peptide interactions.[27–29] Other lipid types can also generate intrinsic curvatures to the membrane to promote or inhibit membrane fusion.[16, 30–32] However, these experimental and computational studies were conducted with simplified membrane compositions, which may not accurately reflect the real situation.

HIV-1 fusion peptides are reported to switch between *α*-helix and *β*-sheet conformations depending on lipid compositions in multiple experiments. Using high-resolution Nuclear Magnetic Resonance (NMR) spectroscopy, circular dichroism (CD), and Fourier-transform infrared spectroscopy (FTIR), Li and Tamm reported that fusion peptide bound to planar POPC/POPG (4/1 mol%) are predominantly α-helical at lower peptide concentration, whereas it assumes anti-parallel β-sheet structure at higher peptide concentration.[33] Recently, Heller showed that fusion peptides are α-helical in the absence of cholesterol, but transition to a β-sheet structure at higher cholesterol concentration (> 30%), using CD and small-angle neutron scattering (SANS).[4, 8] These experimental results indicate that HIV-1 fusion peptide may undergo a helix-to-sheet transition during the fusion process. However, none of these works have membrane compositions that closely resemble human T-cell membranes, which could potentially lead to misinterpretations.

Molecular dynamics (MD) simulation is a powerful tool to investigate the membrane insertion process of various peptides at the atom level across different lipid compositions, although its effectiveness heavily depends on both the accuracy of the forcefields and the accessible timescales.[34–37] Numerous studies show that direct observations of membrane insertion of peptides with conventional MD simulations can be time-consuming due to the energy barriers of lipid membranes. [34–37] The development of coarse-grained MD forcefields enables spontaneous membrane insertion for some peptides within simulation timescales.[34, 38] Additionally, MD simulations have also been performed to study interactions between the HIV-1 fusion peptide and membranes with simple lipid compositions, showing membrane-associated conformation of the HIV-1 fusion peptide. [7, 11] There is an increasing need for direct observations of the HIV-1 fusion peptide inserted into human T-cell membrane mimics in simulations to investigate the in-situ membrane-activated conformational change of the HIV-1 fusion peptide.

In this work, we conducted multiscale molecular dynamics simulations to investigate interactions between the HIV-1 fusion peptide and complex target host membranes. To mimic real environments for inserting HIV-1 fusion peptides, we constructed a complex membrane to simulate well-mixed MT-4 cell membranes[39]. To mimic the geometrical constraints imposed by the rest of the gp41, we restrained the fusion peptide’s orientation and proximity to the membrane. Moreover, we used a multiscale molecular dynamics simulation protocol to accelerate lipid mixing while enabling conformational change of fusion peptide. We observed the membrane insertion, anchoring, and perturbation caused by the HIV-1 fusion peptide to probe its complex roles in facilitating HIV-1 cell entry.

## 2. Methods

To study the membrane interactions of the HIV-1 fusion peptide, we constructed a simulation system consisting of a human T-cell membrane mimic and a fusion peptide restrained on the top of the membrane. By modeling this system in both all-atom and coarse-grained molecular dynamics simulations, as well as umbrella sampling simulations, we investigated the membrane insertion, anchoring, and lipid perturbation caused by the HIV-1 fusion peptide. The detailed description is as follows:

### Complex Host Membrane Setup

Lipid compositions of cells that could be infected by HIV-1 have been measured for different cell types.[39, 40] In this work, we choose MT-4 cell, cancerous infected T-4 cells, to build cell membranes in simulations, since T-4 cells are from immune systems and highly relevant to HIV-1 infection. The phospholipid composition of MT-4 cells has been measured in experiments.[39] In our simulations, we only consider lipids with molar concentrations larger than 5 mol% to maintain computational efficiency, since lipids with low abundance would not reflect their properties in simulations. As shown in **Table 1**, all the lipid molar ratios are recalculated after neglecting those low-ratio lipids. In addition, lipid polyunsaturations in cell bilayer membranes are considered in this work. We mainly use experimental measurements on human blood cells.[41] For PCs, DPPC and POPC are chosen to represent saturated and unsaturated PCs, respectively. For PCs, we chose a ratio of DPPC to POPC of approximately 3:2. Similarly, we choose POPE/PAPE and DPPEE/DOPEE to represent saturated/unsaturated PEs and pl-PEs in a ratio of 1:1. Our broad lipid composition is based on lipidomics results for MT-4 cells,[39] but the asymmetry data is not available in that work. To account for lipid asymmetry, we used asymmetry data available from a blood cell lipidome study.[41], we placed all PEs, pl-PEs, and PSs in the inner leaflet (IL), while all the SMs and approximately 80% of PCs are in the outer leaflet (OL). Cholesterol is almost equally distributed, with an OL:IL ration of 54:46. The simulated MT-4 cell membrane was constructed using the CHARMM-GUI membrane builder.[42–48]

**Table 1:** Lipid compositions of MT-4 cell membrane in simulations. The images of lipid chemical structures are adopted from CHARMM-GUI membrane builder.[42–48]

| Head groups | Fraction (mol%) | Lipid Abbreviation | Lipid Full Name | Chemical Structure (attached figure) | Outer leaflet (mol%) | Inner leaflet (mol%) | Total (mol%) |
| --- | --- | --- | --- | --- | --- | --- | --- |
| PC | 31.5 | DPPC | 1,2-Dipalmitoyl-sn-glycero-3-phosphocholine | (a) | 29 | 9 | 18.9 |
|  |  | POPC | 1-palmitoyl-2-oleoylphosphatidylcholine | (b) | 21 | 4 | 12.6 |
| Chol | 31.5 | Chol | Cholesterol | (i) | 34 | 29 | 31.5 |
| PE | 12.4 | POPE | 1-Palmitoyl-2-oleoyl-sn-glycero-3-phosphoethanolamine | (c) | 0 | 12 | 6.2 |
|  |  | PAPE | 1-palmitoyl-2-arachidonoyl-sn-glycero-3-phosphoethanolamine | (d) | 0 | 12 | 6.2 |
| pl-PE | 11.6 | DPPEE | Dipalmytoil di-plasmenyl | (e) | 0 | 12 | 5.8 |
|  |  | DOPEE | Dioleoyl di-plasmenyl | (f) | 0 | 12 | 5.8 |
| SM | 7.6 | PSM | N-palmitoyl-D-erythro-sphingosylphosphorylcholine | (g) | 16 | 0 | 7.6 |
| PS | 5.4 | POPS | 1-palmitoyl-2-oleoyl-sn-glycero-3-phospho-L-serine | (h) | 0 | 10 | 5.1 |

### HIV-1 Fusion Peptide

The HIV-1 envelope glycoprotein (**Figure 1**), abbreviated as Env, is critical for HIV-1 entering host cells to initiate infection. It consists of two domains, gp120 and gp41, which bind with the plasma membrane receptor CD4 and co-receptor CCR5 or CXCR4 on susceptible cells.[1, 49] Env induces membrane fusion through a series of conformational changes.[1] The HIV-1 fusion peptide, consisting of 15 to 22 hydrophobic amino acids, is located at the N-terminus of the gp41 subunit.[5, 12, 14, 16, 50] The HIV-1 fusion peptides interact with T-cell membranes, providing the perturbation necessary for membrane fusion.[51] More importantly, it has been recognized as a promising vulnerable site to bind neutralizing antibodies.[14] The sequence of the HIV-1 fusion peptide is highly conserved.[16, 17] Unless otherwise stated, the sequence numbering in this work follows the HIV HXB2 numbering scheme (https://www.hiv.lanl.gov), i.e., position numbers relative to HXB2CG in GenBank. The initial structure of the HIV-1 fusion peptide in simulations is chosen from residue 512-545 in BG505 (PDB: 5i8h),[14] which has the identical sequence as a CH505 structure (PDB: 6uda) and a CH848 structure (PDB: 6um6).[52] BG505 is a clade A HIV-1 strain.[53] CH505 and CH848 viral isolates are both clade C.[54, 55] Note that since HXB2 numbering for the Env protein starts from gp120, residue 512 is the N-terminus of gp41. The complete sequence from residue 512-545 is AVGIG AVFLG FLGAA GSTMG AASMT LTVQA RNLL, which consists of the N-terminal fusion peptide (FP: 512-524), the fusion peptide proximal region (FPPR: 525-535), and part of the heptad repeat region 1 (HR1: 536-545). We choose the fusion peptide region of BG505 (PDB: 5i8h) and CH848(PDB: 6um6) because of its completeness. Although their sequences are same, their structures are slightly different. As shown in **Figure 1**, HR1 parts of both fusion peptides appear to be helical, while FP and FPPR could be either helical in CH848(PDB: 6um6) or coiled in BG505 (PDB: 5i8h). We found the fusion peptide domain of CH848 (PDB: 6um6) turns into disordered coils after being solvated in water. As a result, we chose the fusion peptide of BG505 (PDB: 5i8h) in the following simulations. As mentioned before, during the insertion process, the gp41 subunit likely adapts a long triple-stranded coiled coil extending towards the host membrane.[5, 18] This intermediate state of the HIV-1 Env protein has not been fully resolved in experiments. Still, long helical structures of the gp41 subunit can be speculated from the similar influenza hemagglutinin (HA) structure[56] and bridge distances (~ 11 nm) between HIV-1 viral and cell membrane.[18] To mimic the actual insertion condition in simulations, residues 536 to 545 as part of HR1 are fixed as a straight helix at a certain distance to the cell membrane. The separation distance along *z*-axis between the center-of-mass (COM) of the fixed helix and COM of the bilayer membrane is set to 5 nm estimated from of full gp41 extended helix length and spoke length reported recently.[18]

**Figure 1.**
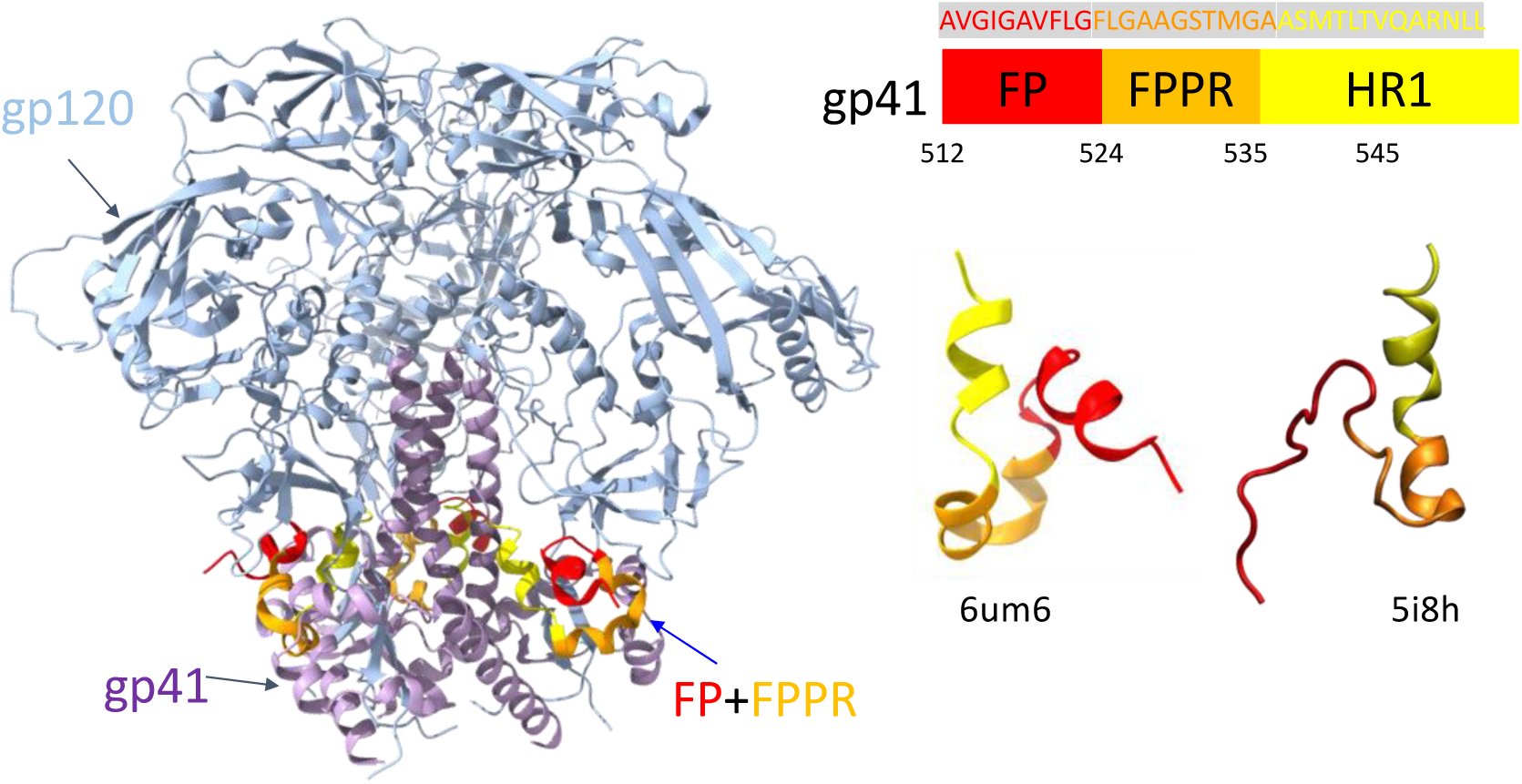
(Left) Structure of HIV-1 trimeric Env protein CH848 (PDB:6um6) and (right) the simulated fusion peptide region from both CH848 (PDB: 6um6) and BG505 (PDB: 5i8h). Both fusion peptides have the same sequences. Other details see Methods.

### Molecular Dynamics Simulations

MD simulations have been recognized as a powerful and high-throughput screening tool to investigate peptide-membrane interactions.[57] Traditional MD simulations calculate forces on every atom to proceed next-step movement, also known as all-atom (AA) MD simulations. As a result, AA MD simulations are limited by their expensive computational cost with larger system sizes. For example, it is time-consuming, especially for the complex membrane in our simulation system, to reach ideal lipid mixing.[58] To increase computational efficiency, many coarse-grained (CG) MD forcefields have been developed by grouping several atoms into CG beads, mostly based on AA simulations in a bottom-up way,[59, 60] i.e., deriving large-scale models from small-scale models. While CG models speed up simulations significantly, the loss of chemical details in CG models often impair quantitative studies of some complex biological processes.[61–63] For example, secondary structures of proteins cannot change during simulations for most CG forcefields due to the lack of hydrogen bond donors/acceptors in CG models.[64] This issue has drawn the attention of CG forcefield developers. Pedersen et al. developed OLIVES, a Go-like model incorporating hydrogen bonding native contacts in the Martini 3 CG forcefield.[65] However, the Go-like model requires a pre-defined set of secondary and tertiary contacts, which we don’t know in advance in our case. Multiscale AA/CG models are developed to combine the high accuracy of AA models and the computational speed of CG models, especially in biological applications.[66–69] Here, we developed a multiscale MD modeling scheme through AA1/CG-AA2 MD simulations as shown in **Figure 2**, which allow us to accelerate lipid mixing and still investigate change of secondary structures of HIV-1 fusion peptide simultaneously. Specifically, in this work, AA simulations are conducted with the CHARMM36 forcefield,[70] which CG simulations are conducted with the Martini 3 forcefield.[62] The CG simulations start at the same configuration of the AA1 initial state obtained from AA-to-CG conversion to examine the influence of multiscale modeling on accelerating membrane insertion and lipid mixing. The AA-to-CG structural conversion of protein and lipids is done through the backward.py[71] script to get the initial input files for the CG simulation. The CG forcefield of the protein is updated by martinize2 and vermouth[72] from Martini forcefield developers.[61–63, 73, 74] One CG water bead is assigned randomly at one Oxygen position for every 4 AA water molecules. Note that, the CG water bead is only assigned for those AA water molecules which have at least 6 neighboring water molecules within 6 Ȧ cut-off radius. This restriction is applied to avoid artificially increasing water density in some low water density area. AA2 simulations start from the final configuration of CG simulations through the CG-to-AA conversion. The CG-to-AA conversion is conducted by the sinceCG.sh[75] script we developed before through a series of energy minimization runs, which is leveraged from backward.py[71]. To recover the original secondary structures in the AA simulations, distance restrains are applied to those residues of the fusion peptide identified by secondary structures other than coils at the end of the last AA run. More details about AA-to-CG and CG-to-AA conversions are shown in the Supplementary Information **Figs. S1 and S2**.

**Figure 2.**
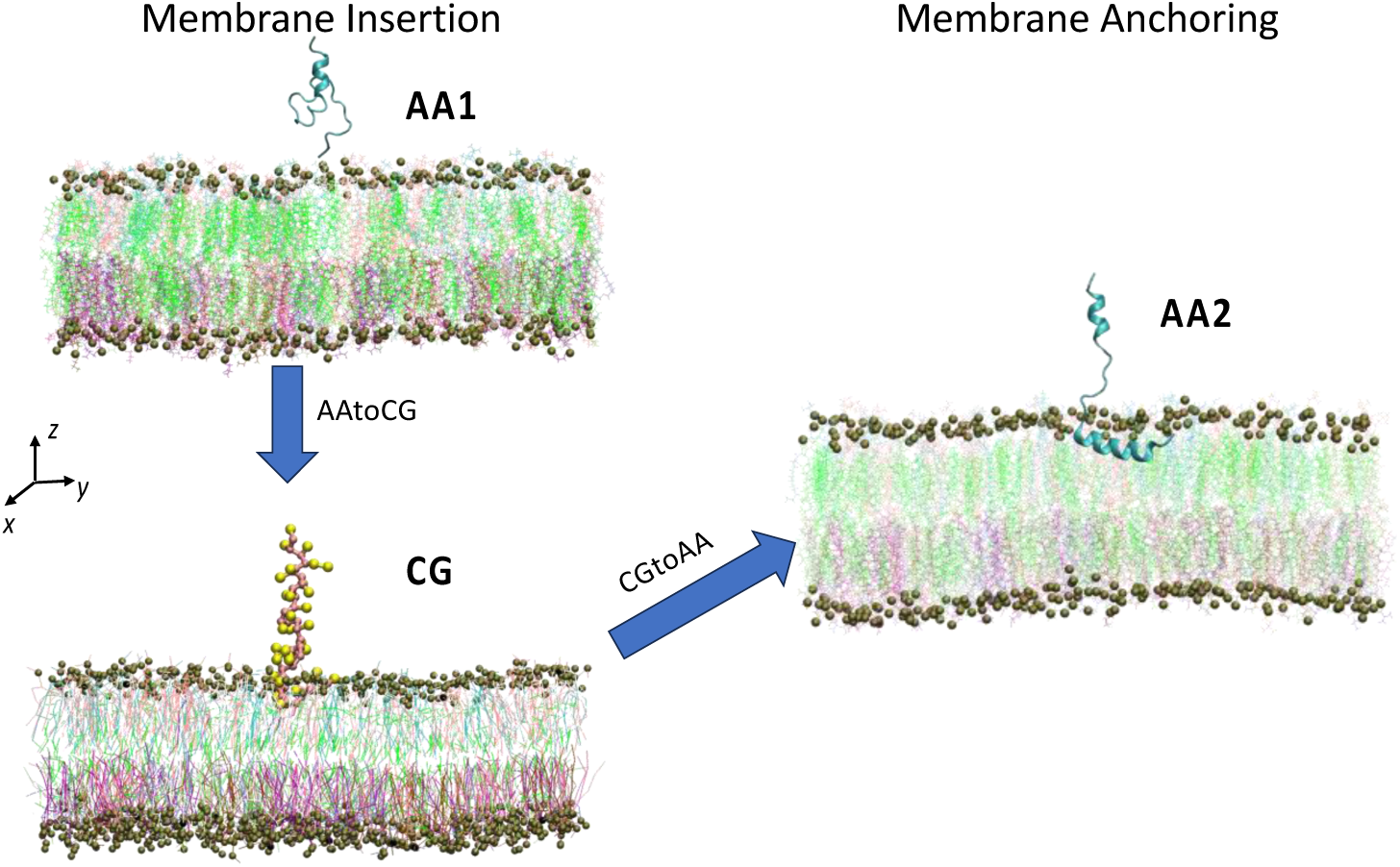
Multiscale simulation systems of MD modeling between AA and CG systems. This multiscale simulation starts with investigations of membrane insertion process in both AA (AA1) and CG systems with HIV-1 FP directly from PDB: 5i8h. After 1 *μs* of AA1 simulations, the FP in the AA1 simulation is only able to either insert with very small portions or briefly touch the membrane. The CG simulation runs very fast with the quick insertion of FP into the membrane in all the independent runs. However, FP cannot stay steadily inside the membrane even after 1 *μs*. Then the CG simulation is backmapped to the AA simulation again (AA2) to investigate the conformational change of the fusion peptide inside the membrane. 5 independent simulations have been conducted for every system, but for each system we only show one snapshot as a representative.

Individual AA and CG simulations follow some general settings as described below. To keep consistency among AA and CG simulations, most simulation setups are the same in both scales unless otherwise stated. Those different settings are mainly due to scale difference between AA and CG. Water was modeled using the TIP3P model. We placed the fusion peptide on top of lipid membrane using Packmol[76] in a 15 × 15 × 15 nm^3^ simulation box with 37 Cl^−^, 86 K^+^, and 79784 AA water molecules (19946 CG water beads) to keep zero net charge in the system. The simulated lipid bilayer membrane is constructed with 315 cholesterol, 190 DPPCs, 125 POPCs, 80 PSMs, 50 POPSs, 60 DOPEEs, 60 DPPEEs, 60 PAPEs, and 60 POPEs molecules, following corresponding distributions in **Table 1**. The forcefield files of AA simulations were generated using CHARMM-GUI.[43, 44] Both AA and CG MD simulations were performed in GROMACS 2022.4.[77, 78] Forces in excess of 1000 kJ/(mol·nm) in the initial state of the system were removed through steepest descent energy minimization. Initial atom velocities were assigned from a Maxwell–Boltzmann distribution at 310 K. Periodic boundary conditions were applied in all three dimensions. Unless otherwise stated, production runs of 1 μs or more in the NPT ensemble at 310 K and 1 bar employing a V-rescale thermostat[79] with a time constant of 1.0 ps and semiisotropic C-resale barostat along *x*-/*y*-axis with a time constant of 4.0 ps (AA) or 12.0 ps (CG) and compressibility of 4.5 × 10^−5^ bar^−1^ (AA) or 3.0 × 10^−4^ bar^−1^ (CG). The equations of motion were numerically integrated using a leapfrog algorithm with a 2 fs (AA) or 10 fs (CG) time step. Lennard-Jones interactions were shifted smoothly to zero at a 1.2 nm cutoff. Electrostatics were modeled using particle mesh Ewald summation with a real-space cutoff of 1.2 nm in AA and using reaction field summation with a real-space cutoff of 1.1 nm in CG. Simulation snapshots were saved for analysis at every 100 ps. Simulation trajectories were visualized using VMD.[80] The COM pulling along *z*-axis direction is applied to restrain the 5 nm distance between COM of FP residue 536 to 545 and COM of the whole bilayer membrane with an elastic constant of 1000 kJ/(nm^2^*mol). Unless specified otherwise, each simulation in this study has been carried out for 5 independent runs with the same settings for statistical analysis. All the independent runs reach 1000 ns (1 μs) simulation time. Analysis in this work was conducted through either GROMACS tools[77, 78] or the MDAnalysis toolkit.[81, 82] The computation speeds are about 55 ns/day for the AA system and 2085 ns/day for the CG system on 1 NVIDIA 4×A100 GPU node.

### Umbrella Sampling Method

We performed two umbrella sampling simulations to obtain free energies when the fusion peptide enters and exits the membrane, respectively. The entering and exiting potential of mean force (PMF) profiles were obtained via umbrella sampling simulations within atomistic settings, respectively. We used distances from the center-of-mass for the whole membrane to the alpha carbons of residues 517 to 521 for entering PMF as the reaction coordinate. For exiting PMF, we use distance from the alpha carbons of residue 529 to 531 (residue 532 to 545 were removed for computational efficiency) and Oxygen (O3) of cholesterols in the lower leaflet as the reaction coordinate. Initially, the entering or exiting pathway was generated through a steered MD simulation starting from disordered fusion peptide outside of the membrane (initial configuration of AA1) or helical fusion peptide inside the membrane, where the fusion peptide was dragged into or outside the membrane using a harmonic potential with a force constant of 1,000 kJ/ (mol·nm^2^) and a rate of 5×10^−4^ nm/ps. For entering PMFs, a total of 20 windows between 2 and 4.3 nm from the membrane core were used, with a spacing of 0.1 nm for the windows from 2 to 3.5 nm, and 0.2 nm for those between 3.5 to 4.3 nm. The separated umbrella simulations were carried out using force constants of 1,000 and 500 kJ/ (mol·nm^2^) for the small and large windows, respectively. For exiting PMFs, we have 20 windows with a space of from 0.2 nm for the distance to the membrane surface from 0.5 nm to 4.5 nm. The first 10 ns of simulations were considered equilibration, and thus excluded from the free energy calculations. The simulation time for each window is about 500 ns for entering PMF and about 300 ns for exiting PMF. The final results of both simulations were obtained using the weighted histogram analysis method (WHAM)[83] with the autocorrelation correction through the GROMACS[77, 78] command “*gmx wham*”[84] and the error was estimated using 100 bootstraps.

## 3. Results and Discussion

### 3.1. Membrane Insertion of HIV-1 Fusion Peptides: AA1 and CG Simulations

In the AA1 simulation, as shown in **Fig. 3a**, HIV-1 fusion peptide with restrained HR1 regions becomes disordered (**Fig. 4**) in the solvent because of highly hydrophobic residues.[51] We did not observe the insertion of fusion peptide into the complex membrane at the end (1000 ns) of this AA1 simulation in **Fig. 3c**. We count the numbers of contacts during this simulation through the GROMACS command “*gmx mindist*” in **Fig. 3e** and others in **Fig. S3**, i.e., numbers of atom pairs between the fusion peptide and the bilayer membrane with a pairwise distance smaller than 6 Ȧ cutoff. The numbers of contacts oscillate several times over the 1 μs AA1 simulation, indicating that the fusion peptide does have many chances to touch the membrane surface, but those contacts don’t create the actual insertion and anchor inside the membrane. Among all 5 independent runs, 1 out of 5 runs (Seed 4) has more frequent protein-membrane contacts, but we found it still cannot insert into the membrane firmly through large oscillations of contacts from and back to 0. The reason behind this behavior could be that there might be an energy barrier for the fusion peptide at the solvent-membrane interface under the current configuration of the AA1 simulations to block it from penetrating the membrane surface. This barrier could also be the penalty for defect formation in the head group salt-bridge network.[85] We also tracked the change of secondary structures as the fusion peptide touches and leaves the membrane in these AA1 simulations in **Fig. 4**. Mostly, when the fusion peptide moves freely in the solvent, the secondary structures are random coils. Interestingly, corresponding to the fusion peptide touching the membrane surface around 500-600 ns, residues 515-521 form *β*-sheet/bridge structures along with turns in Seed 0 (**Fig. 4a**). Especially in Seed 4 (**Fig. 4e**), residue 518-523 form *β*-sheets from 200 ns to 750ns almost continuously corresponding to membrane bounding during that time period (**Fig. S3d**), but those *β*-sheet structures are not stable after the fusion peptide leaves the membrane around 800 ns.

**Figure 3.**
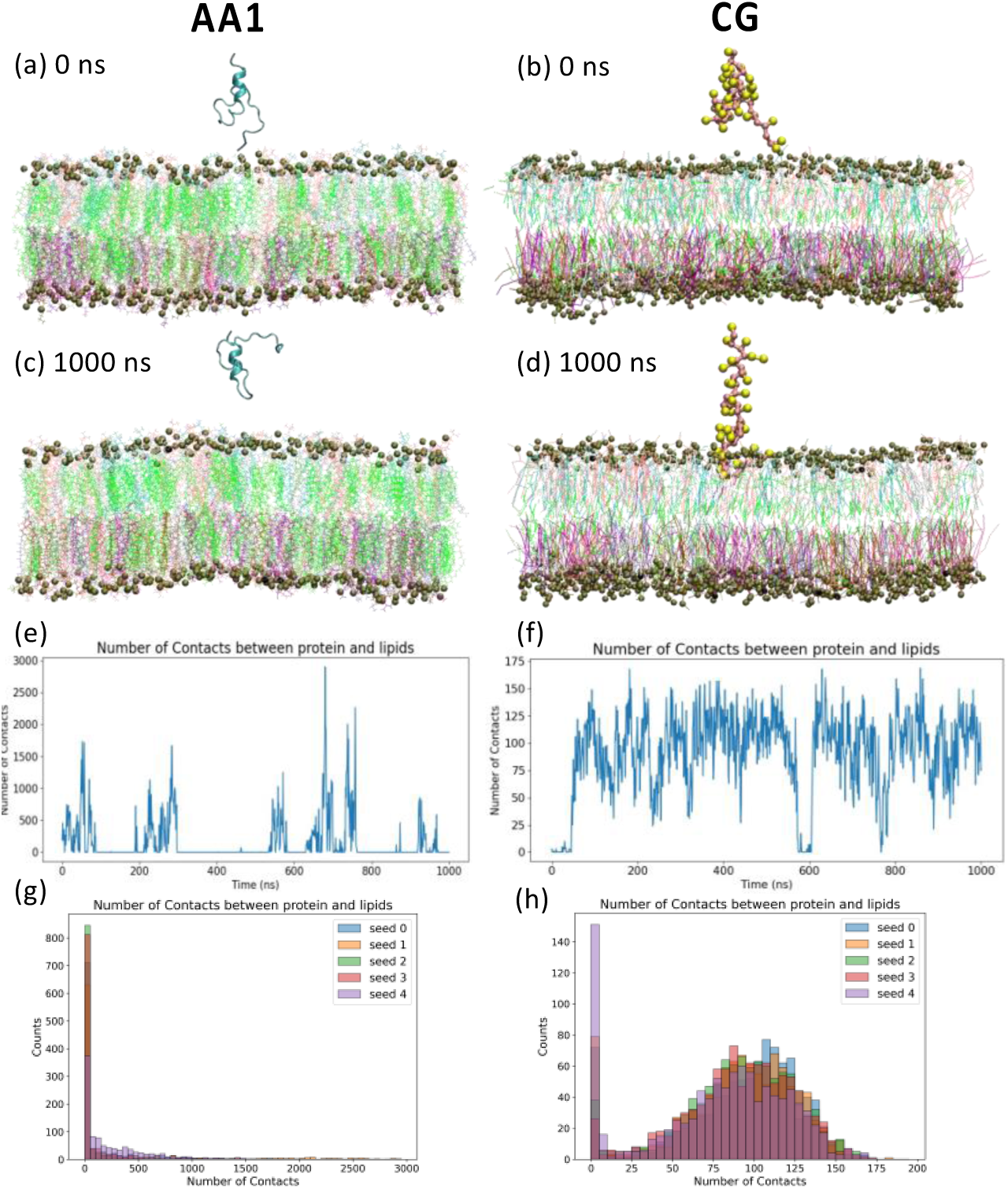
Multiscale modeling of HIV-1 fusion peptides interacting with the complex cell membrane from the same initial configuration, i.e., fusion peptide is completely outside the membrane at ts = 0 ns in the (a) AA1 and (b) CG systems, repectively. The final snapshots are reported at ts = 1000 ns for the (c) AA1 and (d) CG simulations. In those snapshots, the lipids in the membrane are colored based on their types, i.e., cholesterol (green), DPPC (white), POPC (pink), PSM (cyan), POPS (purple), DOPEE (lime), DPPEE (mauve), POPE (ochre), and PAPE (iceblue). Phosphate atoms, and corresponding CG beads, are amplified as dark green balls to represent boundaries of the membrane. Secondary structures of each residue in the fusion peptide are shown in ribbon cartoons at corresponding time steps in the AA1 simulation. Time evolution for numbers of (e) atom-pair contacts in the AA1 simulations and (f) CG bead-pair contacts in the CG simulation within 6 Ȧ cutoff between the fusion peptide and membrane. 5 independent simulations have been conducted for both AA1 and CG system, but only one of those simulations are shown here. Results from remaining simulations are reported in **the Supplementary Information figures S3 and S4**. Histograms for numbers of (eg atom-pair contacts in the AA1 simulations and (f) CG bead-pair contacts in the CG simulation within 6 Ȧ cutoff between the fusion peptide and membrane for all 5 independent simulations.

**Figure 4.**
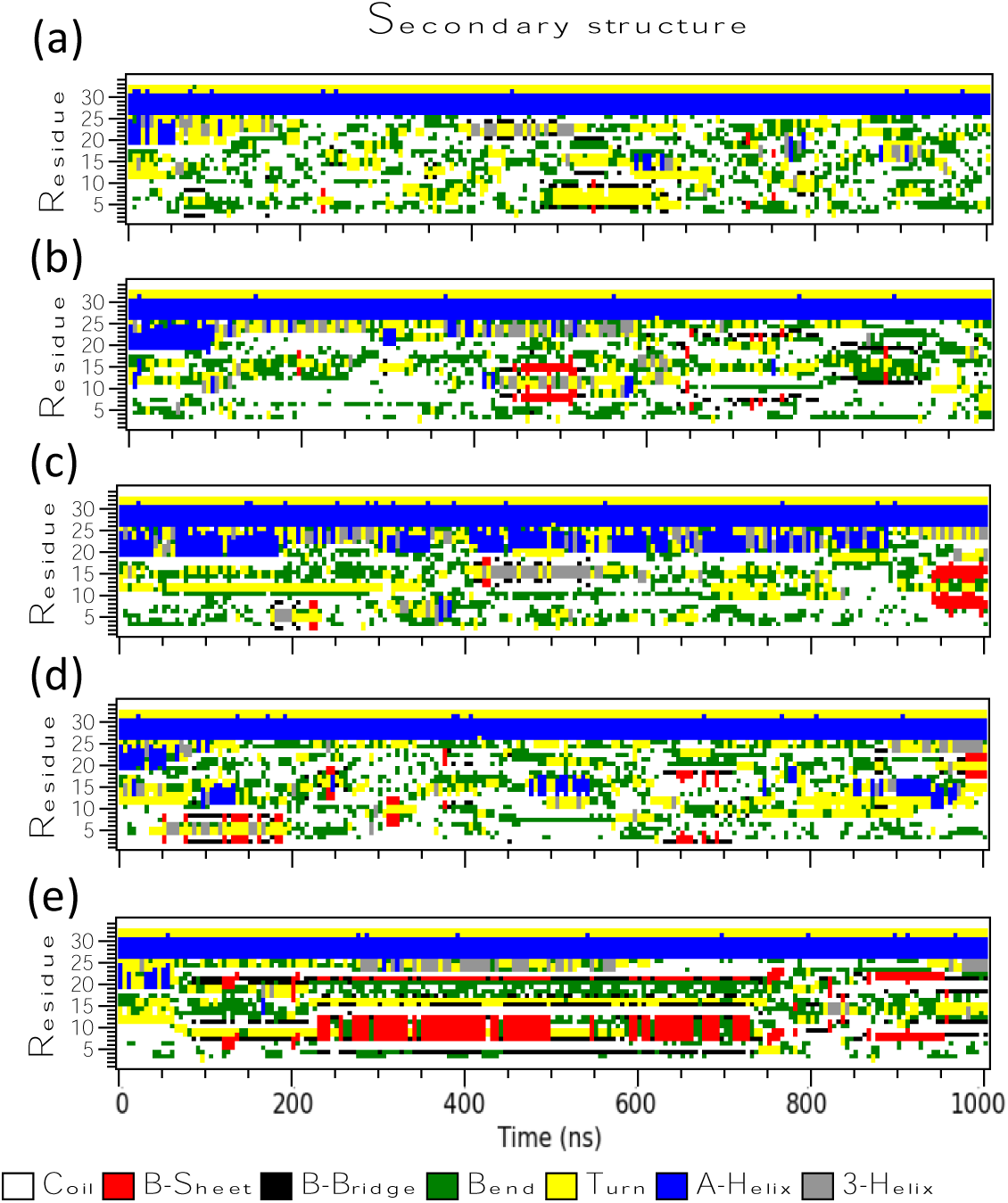
Time evolution for secondary structures of the HIV-1 fusion peptide in the AA1 simulations with 5 independent runs. The secondary structures of the fusion peptides are calculated from the GROMACS command “gmx do_dssp”. Figures are generated from xpm files through the GROMACS command “gmx xpm2ps”.

We examined the existence of this energy barrier through the umbrella sampling method in **Fig. 5**. By pulling the “AXXXG” motif into the membrane, where “XXX” are three continuous hydrophobic residues (VFL in this work), an energy barrier of 16.5 ± 2.9 kJ/mol can be found when the “AXXXG” motif inserts. The membrane boundary is estimated through the average location of proximal phosphorus (P) atoms. The PMF starts to increase when the COM of the pulling group is about 1.5 nm away from the membrane boundary and reaches the highest point at the membrane boundary. Then, the PMF starts to decrease after entering the headgroup region. Though not directly comparable, this energy barrier for membrane insertion of HIV-1 fusion peptide is similar to those of Polytheonamide B (18.0 kJ/mol).[36] We suspect that the reasons for slow membrane insertion in AA simulations could be: (1) membrane insertion of HIV-1 fusion peptide need more than 1 *μ*s of simulations, (2) the HIV-1 fusion membrane insertion requires some specific local environment. Some recent experiments also show that the HIV-1 fusion peptides prefer to insert at the boundary between high and low cholesterol regions, i.e., the boundary between disordered and ordered regions.[9, 22, 29] However, we aim to investigate the insertion ability of HIV-1 fusion peptide under the condition without clear phase separation. Another efficient way to smooth the energy barrier in general is to decrease the system resolution.[86, 87] As a result, we proceed this insertion and anchoring process with lower-resolution simulations, i.e., CG simulations.

**Figure 5.**
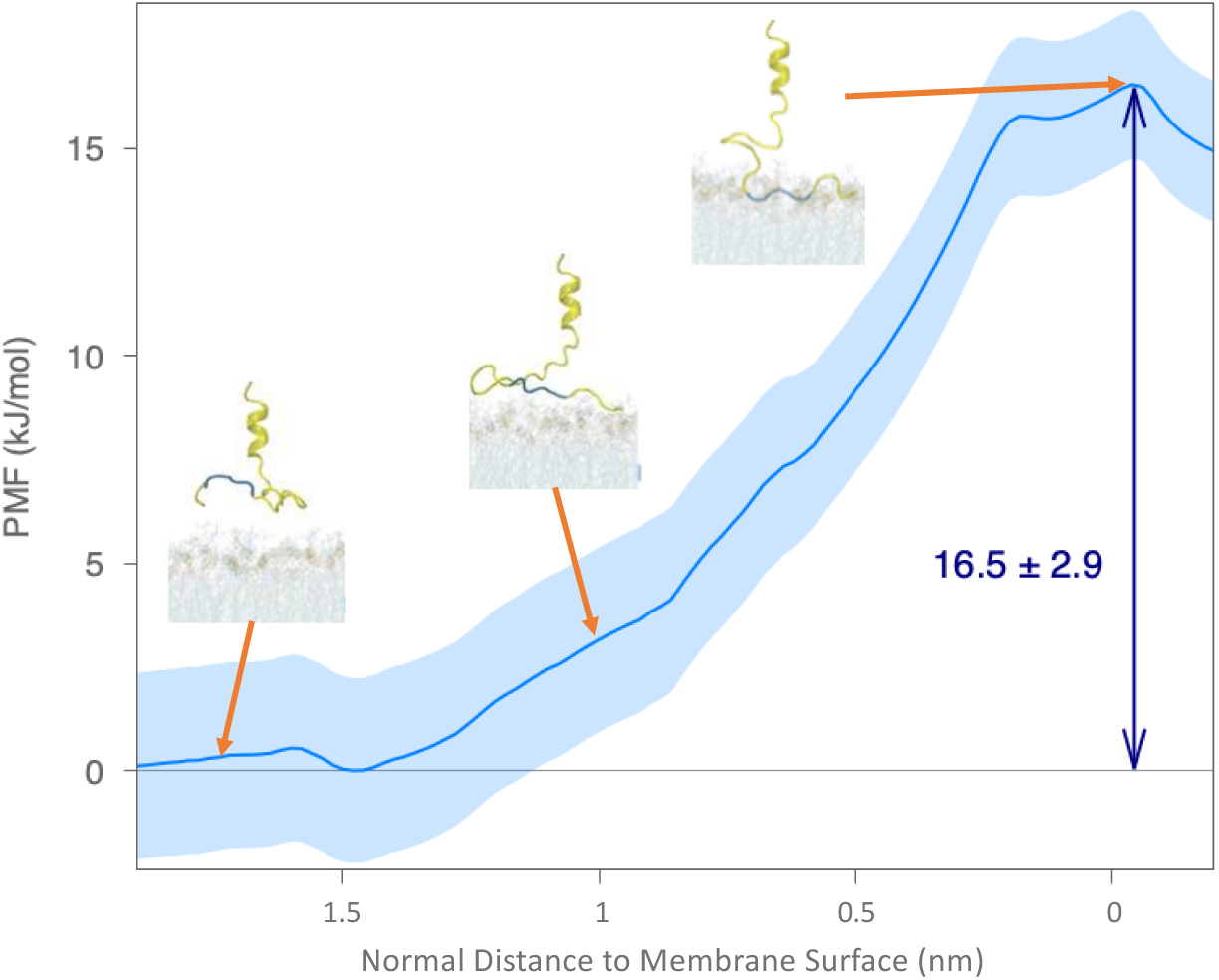
Examination of free energy barriers preventing fusion peptide inserting into the complex membrane in the AA simulations through umbrella sampling methods. Snapshots of fusion peptide and membrane with different insertion depth in membrane are shown with arrows towards approximate points. The PMF profile for the COM of the “AXXXG” motif moving from the solvent into the lipid membrane along the negative z-axis direction. The x-axis represents COM positions of the “AXXXG” motif relative the membrane boundary defined by the average position of the P atoms in the upper leaflet, which is positive in the solvent above the membrane and negative inside the membrane. The PMF curves are computed using umbrella sampling and WHAM, and uncertainties in the light blue shade are estimated by 100 rounds of bootstrap resampling through autocorrelation.

In the CG simulation, the initial configuration of the AA1 system shown in **Fig. 3a** is converted into the coarse-grained scale in **Fig. 3b** through our AA-to-CG conversion shown in **Fig. S1**. We also run CG simulations for 1 *μ*s. The HIV-1 fusion peptide can be inserted into the complex lipid membrane at the end of the CG simulation, as shown in **Fig. 3d**. Time profiles for numbers of contacts in CG simulations, as shown in **Figs. 3f and S4**, indicate that part of the fusion peptide can get into the complex membrane within less than 100 ns. The energy barrier in the CG system is lower at the solvent-membrane interface than in the AA system due to much lower system resolutions and changes in the Hamiltonian. However, the fusion peptide cannot anchor firmly inside the membrane without undergoing a conformational transition. The HIV-1 fusion peptides are assumed to stay firmly inside host membranes to drag host membranes closer to the viral membrane during the gp41 refolding process from the hairpin intermediate[18, 19] to the postfusion state.[1] The conformational switchovers of the fusion peptide during interactions between proteins and membranes are widely recognized to be critical during the HIV-1 cell entry process.[4, 5, 8, 13, 16, 88, 89] Unfortunately, conformational changes of the HIV-1 fusion peptide in the host membrane cannot be captured in the CG since the Martini 3 CG forcefield used in this work doesn’t account for secondary structural changes. Therefore, we converted the final snapshot of the 1 *μ*s CG simulation into an all-atom (AA2) system to study membrane anchoring behaviors of HIV-1 fusion peptides (see next section).

Interactions between membranes and membrane-active peptides are also dependent on lipid environments.[90] To investigate lipid dynamics across different simulation scales, we calculated lipid mixing and protein-lipid contacts in AA1 and CG simulations, respectively. The lateral radial distribution function of different lipids with respect to the fixed helix are calculated through the GROMACS command “*gmx rdf*” in the early stage (0-50 ns) and late stage (950-1000 ns) shown in **Figs. 6**. Although lateral radial distribution functions in both AA1 and CG systems exhibit clear peaks at the early stage indicating incomplete lipid mixing, those profiles of all four lipids in the CG system approach flat to 1 at the late stage, contrasting with big fluctuations observed in the AA1 system. Other lateral radial distribution functions are reported in the Supplementary Information **Fig. S5** for AA1 simulations and **Fig. S6** for CG simulations, respectively. Consistently, all other independent AA1 simulations have the similar lipid mixing behavior as shown in **Figs. 6a and 6c**. Comparing to AA runs, lateral radial distribution functions of other CG simulations are similar in the early stage but different in the late stage. However, these CG runs tend to remain flat over long-distance ranges (r > 4 nm) yet exhibit instability and inconsistency in shorter distance ranges, suggesting that prolonged insertion of the fusion peptide in CG simulations may significantly disturb nearby lipid distributions Our results suggest that rapid dynamics in CG simulations allow faster lipid mixing than AA1 simulation, even under the influence of inserted fusion peptide.

**Figure 6.**
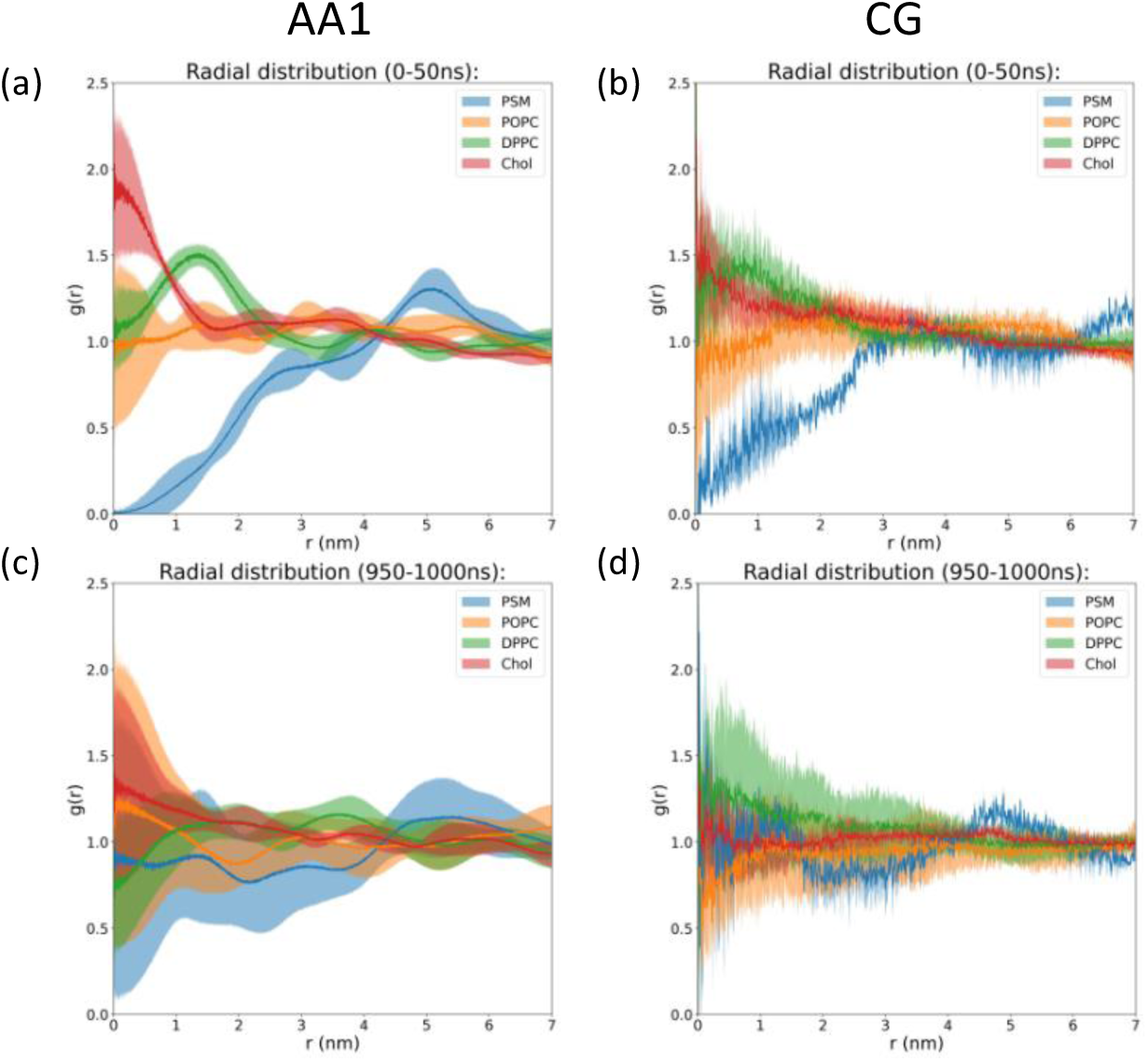
Radial distribution functions of projection from the fixed helix on surface and lipids in the outer leaflet at the (a, b) early stage (0-50 ns) and (c, d) late stage (950-1000 ns) for the (a, c) AA1 and (b, d) CG systems, respectively. Notably, these calculations of radial distribution functions use only x and y components of the distance, i.e., they are 2-dimentional lateral radial distribution functions along the complex membrane. Only Chol (cholesterol), DPPC, POPC, and PSM are calculated here since they are lipids in the upper leaflet. The bin sizes are 0.002 nm for AA1 systems and 0.01 nm for CG systems. The shaded areas represent standard deviations calculated from 5 independent simulations for both AA1 and CG system. Results from individual simulations are reported in the Supplementary Information **figures S5 and S6**.

To compare protein-lipid interactions during membrane insertion across different simulation scales, we counted protein-lipid contacts for different lipid types in AA1 and CG simulations, shown in **Figs. 7, S7, and S8**. Notably, POPC and DPPC lipids are in contact with fusion peptide when it first touches the membrane in both AA1 and CG simulations. DPPC lipids have slightly higher numbers of contacts in AA1 simulations than POPC, especially for those AA1 simulations without long-time insertion, comparing to nearly equivalent numbers in CG simulations. Another difference in protein-lipid contacts between AA1 and CG simulations is the contact between fusion peptide and cholesterol. There are almost no protein-cholesterol contacts reported in AA1 simulations, but all CG simulations see stable contacts between fusion peptide and cholesterol. Since cholesterol molecules are smaller and located near the membrane center, the difference in protein-cholesterol contacts also indicates the deeper insertion in CG simulations. Interestingly, as shown in **Fig. 7d**, CG simulations have a consistent pattern for histograms for numbers of contacts between fusion peptides and different lipids across 5 independent runs, but we cannot observe a similar trend in AA1 simulations in **Fig. 7c**, likely due to different insertion behaviors of fusion peptides in AA1 and CG simulations. A number of contacts among the same lipid types can also show the completeness of lipid mixing, as shown in **Figs. 7, S7, and S8**. Similar to those reported in **Fig. 6**, oscillation in AA1 simulations indicates incomplete lipid mixing, contrasting with quick mixing in CG simulations.

**Figure 7.**
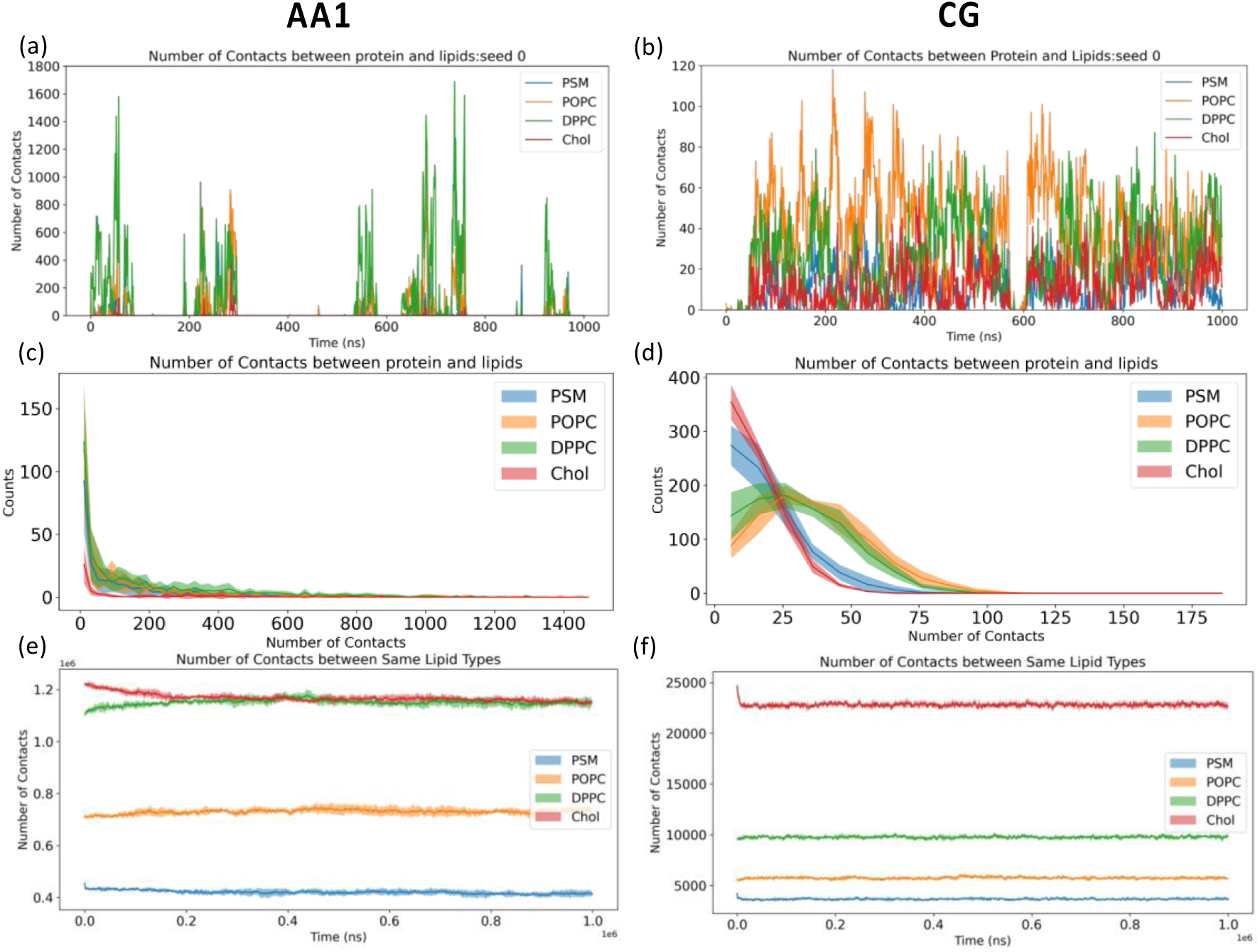
(a, b) Time evolution for number of contacts between protein and lipids in the upper leaflet, (c, d) histograms for nonzero numbers of contacts between protein and lipids in the upper leaflet, (e, f) time evolutions for numbers of contacts among the same types of lipids in the (a, c, e) AA1 and (b, d, f) CG system. Only Chol (cholesterol), DPPC, POPC, and PSM are calculated here since they are lipids in the upper leaflet. 5 independent simulations have been conducted for both AA1 and CG system. The shade areas in (c-f) represent standard deviations calculated from 5 independent runs. Results from all the individual simulations are reported in the Supplementary Information **figures S7 and S8**.

### 3.2. Membrane Anchoring of Fusion Peptides: AA2 Simulation

We convert the final configuration of the CG simulation to the starting configuration of AA2 simulations through our CG-to-AA conversion reported in Supplementary Information **Fig. S2**. AA2 simulations were started with the converted final CG configuration where the fusion peptide inserted inside the host membrane, and simulations were continued for 1 *μ*s. The start and final snapshots of this AA2 simulation are shown in **Fig. 8a**. We calculated the numbers of contacts for protein and membrane in **Figs. 8b and S9**, where steady non-zero contacts between fusion peptide and lipids indicate firm membrane anchoring in the AA2 simulation. However, in 1 out of 5 runs (seed 2), the fusion peptide leaves the membrane at around 480 ns. To understand the reasons for this behavior, we also examine the change of secondary structures during this process shown in **Figs. 8c and S10**. Interestingly, in those AA2 simulations with fusion peptide anchoring inside the membrane, the fusion peptide always forms some helices. while some even gradually grow into a longer helix. In the AA2 simulation reported in **Fig. 8c** (seed 0), although it starts with disordered coils, the fusion peptide in the complex membrane initiates helical folding in a few nanoseconds. The folding process of the fusion peptide can be captured in the traditional identification of secondary structures based on geometrical relationship (**Fig. 8c**) and machine-learning analysis of time-lagged independent component analysis (tiCA) in **Fig. S11**. We also noticed that the growth of helical parts varies in other independent runs, indicating that the ability to form stable long helix might be sensitive to the local lipid environment.

**Figure 8.**
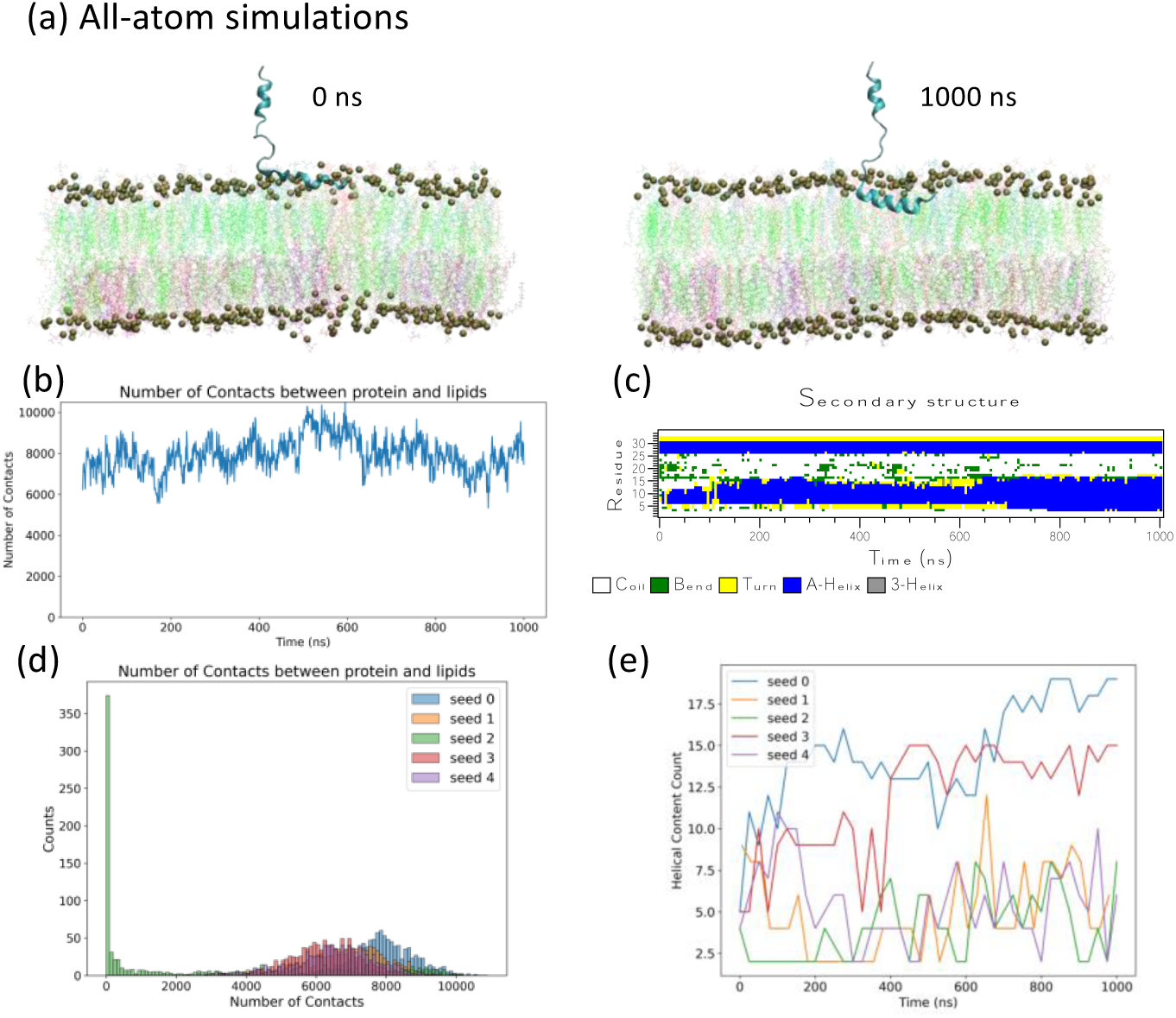
Fusion peptides fold into long helices and lay horizontally in the upper leaflet of cell membrane in the second all-atom simulations backmapped from the coarse-grained simulations. (a) A snapshot of the all-atom simulation system after the fusion peptide inserts into the membrane at the beginning (*t_s_* = 0 ns) and the end (*t_s_* = 1000 ns) of the simulations, respectively. The lipids in the membrane are colored based on their types same as those in Fig.3. Phosphate atoms are amplified as dark green balls to represent boundaries of the membrane. (b) Time evolution for numbers of atom-pair contacts in the AA2 within 6 Ȧ cutoff between the fusion peptide and membrane. (c) Time evolution of the secondary structures for each residue during the 1 *μ*s simulation course. 5 independent simulations have been conducted for the AA2 system, but only one of those simulations are shown in (a-c). Time evolutions for numbers of contacts between protein and lipids from remaining simulations are reported in the Supplementary Information **figures S9**. Time evolution of secondary structures calculated from remaining simulations are reported in the Supplementary Information **figures S10**. (d) Histograms for numbers of contacts between protein and lipids from all 5 independent simulations. Note that fusion peptide in Seed 2 leaves the membrane around 480 ns. (e) Time evolutions for numbers of helical residues from all 5 independent simulations, calculated from secondary structure results reported in (c) and **Figure S10**.

Throughout the AA2 simulation with the anchored fusion peptide, helices inside the membrane lay horizontally in the upper leaflet. The horizontal orientation of the long helix has a larger lateral cross-section area than the vertical orientation, given the amphipathic character of the amino acids of the fusion peptide, which provides a stronger mechanical strength to anchor inside the membrane. Moreover, the numbers of contacts between fusion peptides and membrane are summarized as histograms shown in **Fig. 8d**. Except Seed 2 which leaves the membrane at around 480 ns, fusion peptides in other simulations have relatively stable numbers of contacts with membrane around 6000-8000 atom pairs, which is much bigger than those in AA1 simulations, indicating deeper insertion in AA2 simulations. The helical contents, i.e., numbers of helical residues, from all 5 simulations are also summarized in **Fig. 8e**. The numbers of helical residues are calculated from the sum of *α*-helix and 3-helix residues in **Figs. 8c and S10**. 2 out of 5 independent simulations (Seed 0 and Seed 3) form long helices gradually over the course of simulations.

The reasons why the monomer HIV-1 fusion peptide forms a horizontal helix inside complex membranes are studied from two aspects. First, we examined the structural details of the helix formed by HIV-1 fusion peptide in simulations. The helix wheel diagram in **Fig. 9** is plotted from the membrane-bound helical portion at 1000 ns of seed 0. The sequence of the HIV-1 fusion peptide simulated in this work consists of alternating hydrophobic and neutral residues, i.e., an amphipathic-like helix. As shown in **Fig. 9**, the spatial distribution of those helical residues shows neutral residues close to the membrane surface and hydrophobic residues deeply embedded inside the membrane. This spatial distribution of residues explains the horizontal orientation inside the membrane in simulations. Second, we conducted additional enhanced sampling simulations to study the energy required to pull a helical fusion peptide out of the membrane, as shown in **Fig. 10**. The simulations consist of umbrella sampling windows starting from the final configuration of Seed 0, where a long horizontal helix forms, and ending when the fusion peptide leaves the membrane completely. We observe the helix fall apart as the fusion peptide departs the membrane. The overall energy barrier in this simulation is 77.9 ± 1.5 kJ/mol, which is much higher (61.4 kJ/mol) than it is when pulling the fusion peptide into the membrane in **Fig. 5**. The difference between exiting and entering energy barriers is due to the formation of the helix. In other words, the helix formation enhances the anchoring strength of the fusion peptide inside the membrane. The formation of a horizontal long helix inside the membrane comes from its sequential and structural features, which help the fusion peptide as a membrane anchor.

**Figure 9.**
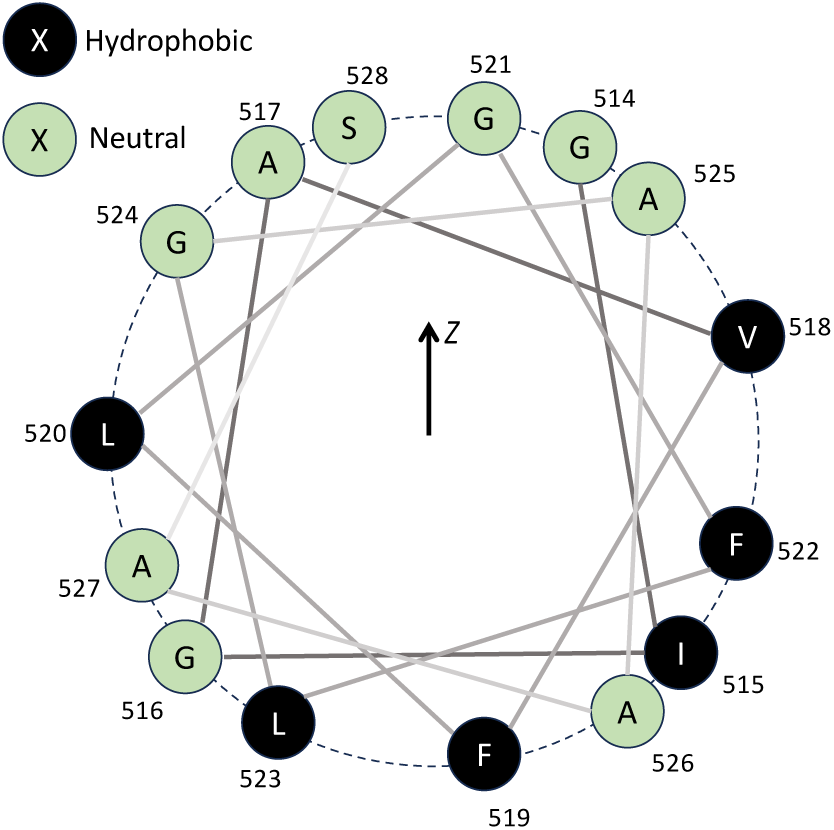
Helical wheel diagram of HIV-1 fusion peptide with helical residues (residue 514-528) at 1000 ns of Seed 0 in simulations. The residue positions in the wheel also reflect their relative locations along z-axis in the system. The green circles are neutral residues, while the black ones are hydrophobic.

**Figure 10.**
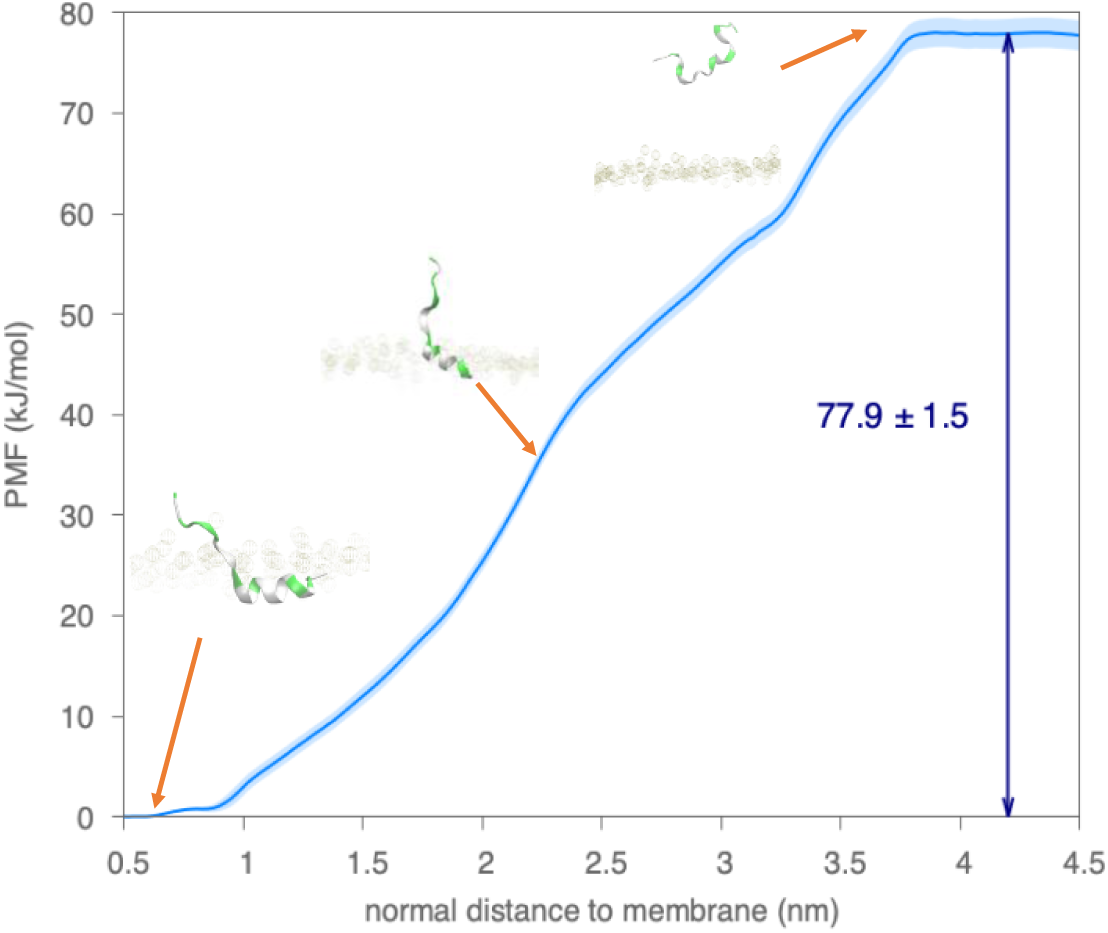
Examination of free energy barriers preventing helical fusion peptide getting out of the complex membrane in the AA simulations through umbrella sampling methods. Snapshots of fusion peptide and membrane along the path are shown with arrows towards approximate points. The PMF profile for the COM of residue 529 to 531 moving into the solvent along the positive z-axis direction. The x-axis represents COM positions of residue 529 to 531 relative the membrane boundary defined by the average position of the P atoms in the upper leaflet, which is positive in the solvent above the membrane and negative inside the membrane. The PMF curves are computed using umbrella sampling and WHAM, and uncertainties in the light blue shade are estimated by 100 rounds of bootstrap resampling through autocorrelation.

Next, we examined the role of fusion peptide in perturbing the lipid environment to promote membrane fusion. Since the complex membrane in our simulations is a close mimic of the human plasma membrane, we can probe *in-situ* scenarios for fusion-peptide-induced lipid reorganization. We examined the near-protein lipid environment first by computing the numbers of contacts between fusion peptides and lipids in the upper leaflet, reported in **Figs. 11a and S12**. In all the AA2 simulations, we observed increased POPC contacts and decreased PSM contacts of the fusion peptide. We observed decreased contacts with cholesterol for those simulations forming long helices (Seed 0 and Seed 3). We counted the lipid distribution within the 2 nm cut-off radius of every membrane-bound amino acid (residue 512-529) at the early stage (0-50 ns) and late stage (950-1000 ns) shown in **Figs. 11 and S13**. Initially, the lipid composition around the fusion peptide was enriched in cholesterol and PSM. At the late stage, cholesterol and PSM density decreased around the fusion peptide. The cholesterol depletion could indicate a change in membrane organization towards a disordered local domain.

**Figure 11.**
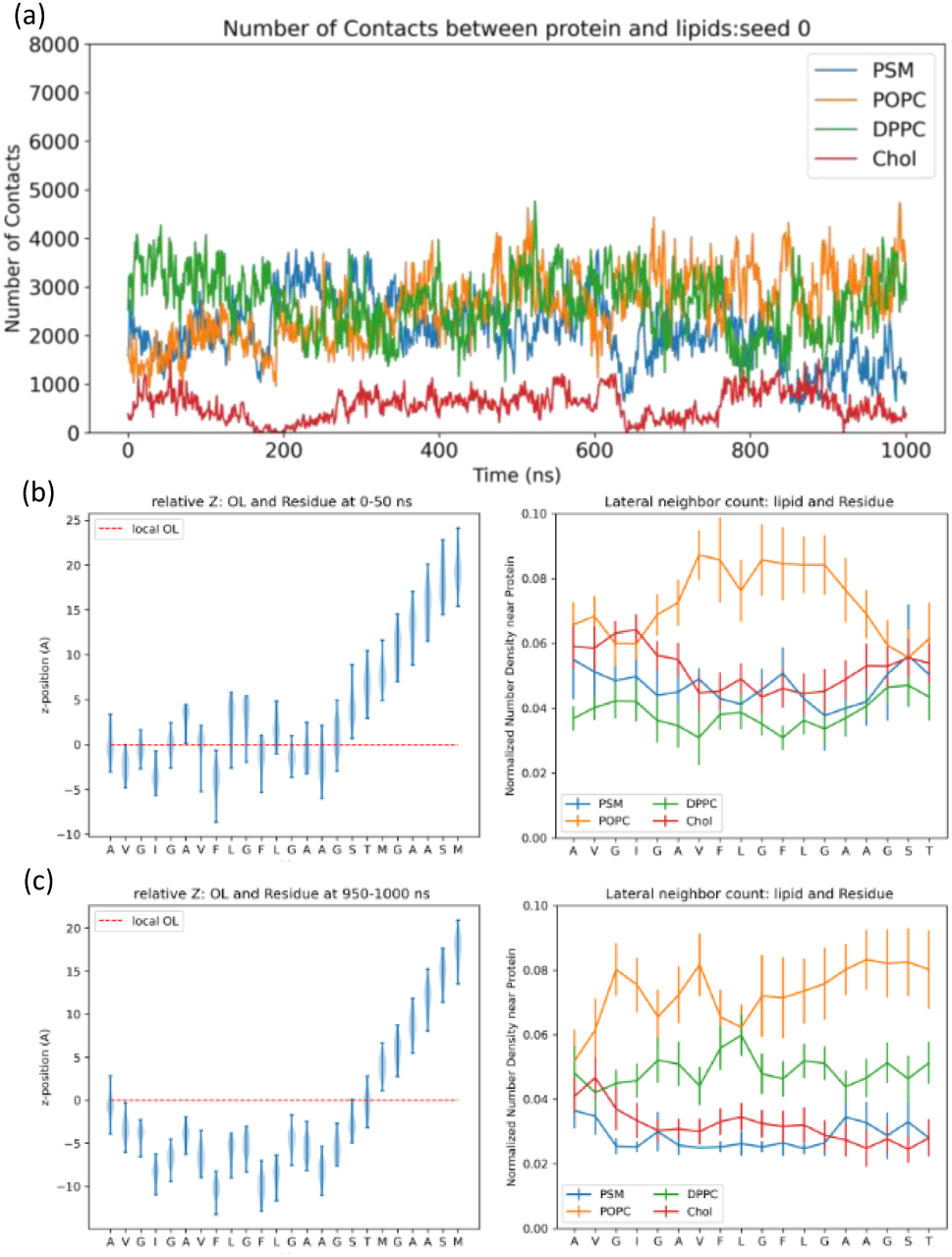
Insertion depth and lipid organization after the HIV-1 fusion peptide inserted inside the membrane. (a) Time evolution for numbers of contacts between fusion peptide and different lipids within 6 Ȧ cutoff. Average insertion depth and nearby lipid distribution of each residue of HIV-1 fusion peptide collected from (b) the early stage (0-50 ns) simulation and (c) late stage (950-1000 ns) simulation. The x-axis coordinates represent residues starting from the N-terminus (residue 512). The y-axis coordinates are the relative distances along z-direction between the COM position for each residue and extrapolated P-atom z-position on the residue COM location representing the boundary of upper leaflet. The x-axis represents each amino acid in the fusion peptide. The y-axis is the lateral number density of certain lipids in the upper leaflet within 2.0 nm cutoff radius normalized by the total number of the corresponding lipid type in the upper leaflet. The lipid number densities are the average values calculated from 50 frames. The error bars represent the standard deviation calculated from 50 frames as well. 5 independent simulations have been conducted for the AA2 system, but only one of those simulations are shown here. Results from remaining simulations are reported in the Supplementary Information **figures S12 and S13**.

Next, we considered more global (non-local) changes in the membrane environment upon inserting the fusion peptide. The density maps of lipids in the upper leaflet at the early-stage (0-50 ns) and late-stage (950-1000 ns) are shown in **Figs. 12 and S14**. First, when fusion peptides stay inside the membrane (except Seed 2), all the lipid types form more significant aggregation in the late stage compared to the early stage. Second, when long helices form (Seed 0 and Seed 3), we found a decreased number density of cholesterol and PSM around fusion peptides in the late stage, which is consistent with observations in **Fig. 11**. The deepest insertion depth of the fusion peptide was only about 1 nm. As shown in **Figs. 11 and S13**, insertion depth profiles show that the fusion peptide does not go deeper inside the membrane but stays near the headgroup region and forms a helix. This is energetically favorable as it satisfies the amphipathic character of the fusion peptide as shown in **Fig. 9**.

**Figure 12.**
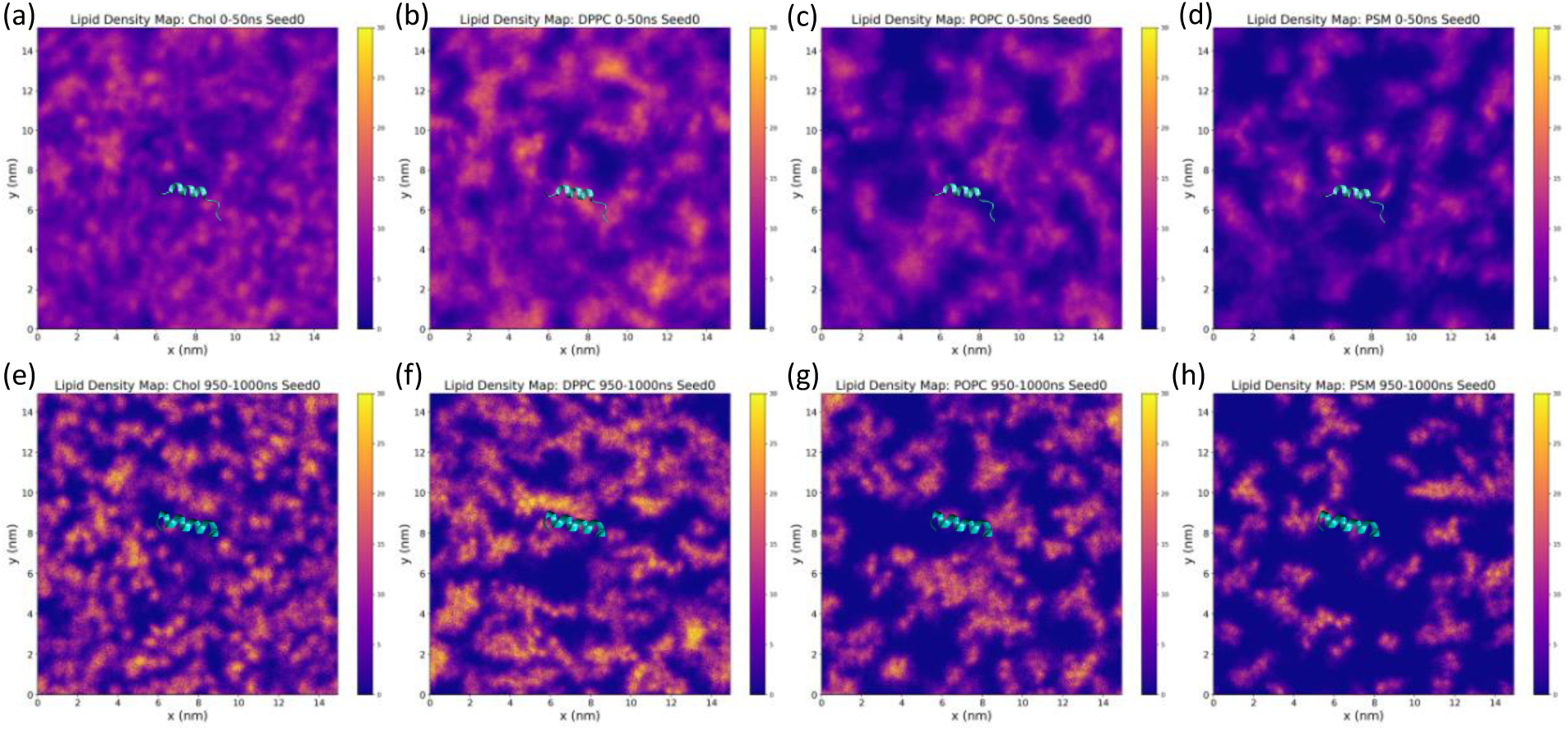
Average number density maps of lipids on the upper leaflet at (a-d) the early stage (0-50 ns) and (e-h) the late stage (950 - 1000 ns). The membrane-embedded portions (residue 512 to 529) of the HIV-1 fusion peptide at 25 ns for the early stage and 975 ns for the late stage are shown in cyan ribbons. The color bars have the same scale from 0 to 30 for all the plots. These density maps were computed from the GROMACS command “gmx densmap”. 5 independent simulations have been conducted for the AA2 system, but only one of those simulations (seed 0) are shown here. Results from remaining simulations are reported in the Supplementary Information **figures S14**.

Finally, we explored whether there is a conserved sequence motif within the fusion peptide that could initiate the folding of the helix. As shown in Fig. 13a, helical structures form around residues 517-521 within several hundred picoseconds. Residues 517-521 are the “AXXXG” motif with “XXX” as three less conversed and hydrophobic amino acids in Fig. 13b. In most AA2 simulations, as marked in Fig. S11, the “AXXXG” motif was always among those regions where the folding of the helix started. Coincidently, the “AXXXG” motif has drawn the attention of structural biologists studying viral membrane fusion. A recent study found that an “AXXXG” motif in the fusion peptide region of SARS-CoV-2 Spike protein is critical in forming the blunted cone shape inside the membrane during the postfusion state.[91] Moreover, from Fig. 13b, one can realize that despite alternating hydrophobic and neutral residues in the HIV-1 fusion peptide region, three Xs in the “AXXXG” motif are the only part that consists of three consecutive hydrophobic amino acids. This amphipathic nature enhanced by the “AXXXG” motif is likely why the helix starts folding from this region. We believe this “AXXXG” region could be one of the critical motifs in HIV-1 fusion peptide that could modulate fusogenic activity.

**Figure 13.**
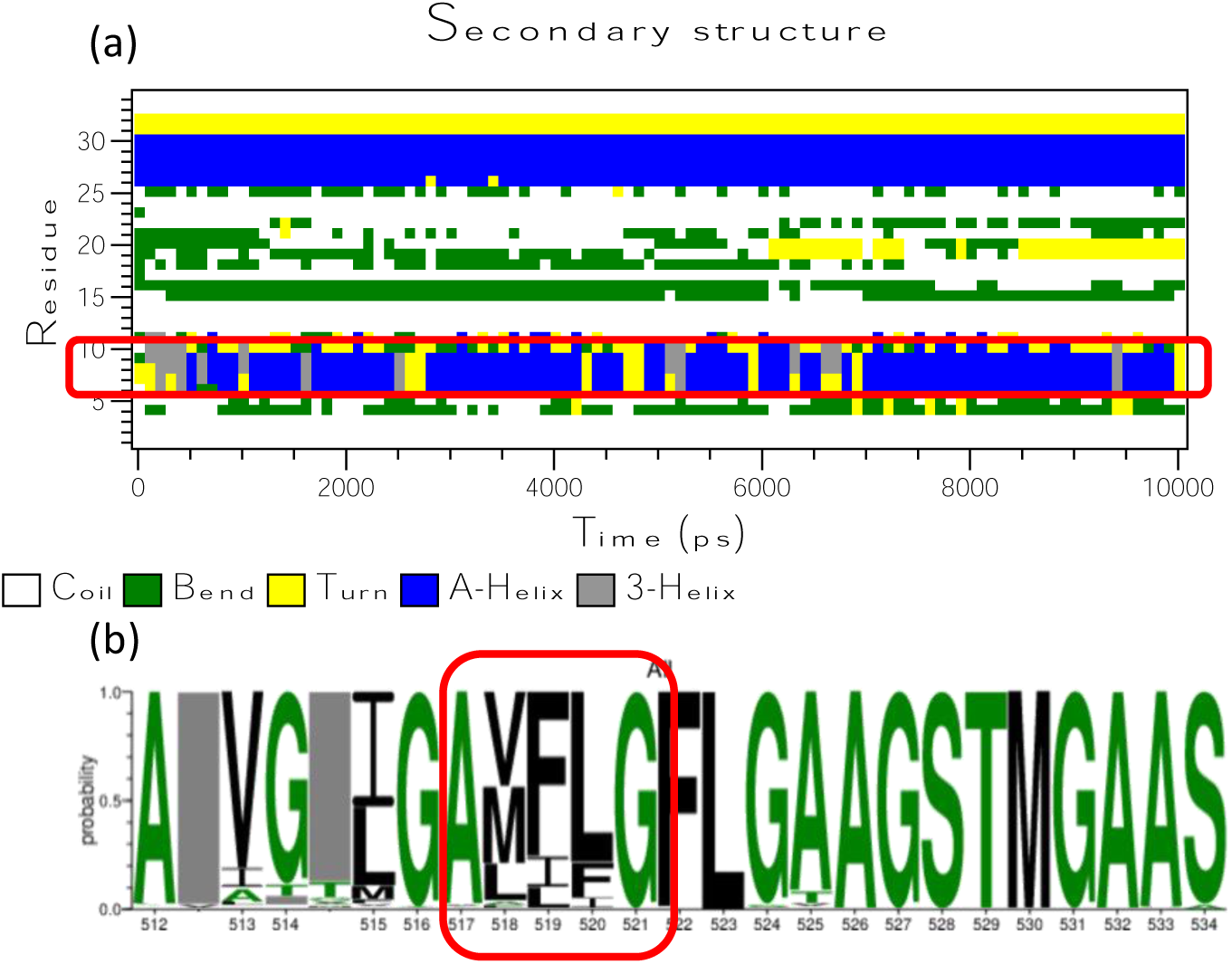
Initial folding “AXXXG” motif of HIV-1 fusion peptide. (a) The secondary structures of each residue during the first 10 ns after the fusion peptide inserts inside the membrane. As circled by the red rectangle, the folding of the helix starts from the “AXXXG” motif. 5 independent simulations have been conducted for the AA2 system, but only one of those simulations are shown here. Results from remaining simulations are reported in the Supplementary Information **figures S11**. (b) Sequence conservation of the HIV-1 fusion peptide region (residue 512 to 534) is investigated among 6481 sequences from LANL HIV database. The HIV-1 fusion peptide region is highly conserved comparing to other domains of the Env protein. The sizes of the letters correspond to the probability of a certain amino acid appearing at the certain position. The character colors represent its hydrophobicity. The green ones are neutral residues, including SGHTAP amino acids. The black ones are hydrophobic residues, including RKDENQ amino acids. The rest amino acids are hydrophilic RKDENQ residues, which are not shown here would be shown in blue.

## 4. Conclusion

A thorough examination of the insertion of the HIV-1 fusion peptide will provide a fundamental understanding of the initial viral-host membrane encounter, which is necessary for potentially developing drugs and vaccines aimed at HIV-1 cell entry. Direct experimental observations of this fusion process at the atomistic level are still challenging. Atomistic (AA) molecular dynamics simulation is a powerful tool to study these fast dynamical processes. Unfortunately, it is still challenging to probe membrane insertion processes with conventional unbiased AA due to varying timescales imposed by the surface tension at the solvent-membrane interface, slow lipid mixing of the complex membrane, and slowly relaxing protein conformational degrees of freedom. For protein-membrane systems, the newly developed MARTINI 3[62] CG forcefield has shown the potential to overcome many of these challenges[61, 92–95] but struggles to accurately update the secondary structures of proteins, which is critical for membrane insertion and anchoring. In this work, we capitalize on the strengths of both AA and CG simulations by implementing an iterative multiscale workflow, AA1 to CG to AA2…, to elucidate the interactions between the HIV-1 fusion peptide and complex T-cell membrane mimic.

In the AA1 simulations, we observe that the fusion peptide makes many contacts with the T-cell membrane but fails to insert. Supporting biased AA simulations indicate that the solvated fusion peptide needs to overcome a large energy barrier for insertion. It is also possible that favorable local membrane changes could facilitate an energetically favorable insertion. The timescale needed to observe these events is much longer than the microsecond timescale considered for AA1 simulations in this study. By generating CG simulations from these AA1 simulations, we overcome the sampling limitations in membrane insertion and lipid mixing. With CG, we observe the insertion of fusion peptide into membranes. Then, we switch back to atomistic (AA2) simulations since the CG couldn’t accommodate the conformational changes in fusion peptide induced by the membrane environment. In AA2 simulations, fusion peptide folds into a helix and lies horizontally in the upper leaflet of the T-cell membrane. Additional biased AA simulations confirm that pulling the folded helix out of the membrane requires much more energy than inserting the fusion peptide into the membrane.

Together, these simulations reveal several molecular insights into the insertion and anchoring mechanisms of HIV-1 fusion peptide. First, even though the fusion peptide is disordered in an aqueous environment, it adopts a helical conformation upon entering the T-cell membrane. Second, this folding is necessary to establish mechanical strength so that the fusion peptide can also function as an anchor to eventually bring the viral and T-cell membranes together. The location and the horizontal orientation of the folded helix imposed by the amphipathic character of the sequence maximize the favorable energetic interactions between the fusion peptide and the complex membrane. Third, the insertion of peptide perturbs the local lipid environment. The reorganization of the lipid implies a disordered membrane that could be favorable for fusion. It should be noted that the above observations were made by considering only a fusion peptide monomer. These behaviors can be altered when considering the fusion peptides from the neighboring protomers from the gp41 trimer. Finally, by considering the sequence conservation profiles of the HIV fusion peptide, we identify the “AXXXG” motif responsible for initiating the helical folding of the fusion peptide from our simulations. Further studies are warranted to delineate the sequence-dependent changes in this motif and the differences between the fusogenic activities of different HIV-1 clades.

## Acknowledgements

This study was supported by the National Institute of Allergy and Infectious Diseases of the National Institutes of Health and by the Duke Center for HIV Structural Biology, grant number U54-AI170752-01. The authors would also like to thank the computational resources provided by the LANL Institutional Computing. This work was performed at the Los Alamos National Laboratory, which is operated by Triad National Security, LLC, for the National Nuclear Security Administration of the U.S. Department of Energy (contract 89233218CNA000001). The views expressed in this article are those of the authors and do not reflect the official policy or position of the U.S. Government.

## Notes

### Competing Interest Statement

The authors have declared no competing interest.

