## Supplemental Files for "Insertion and Anchoring of HIV-1 Fusion Peptide into Complex Membrane Mimicking Human T-cell"

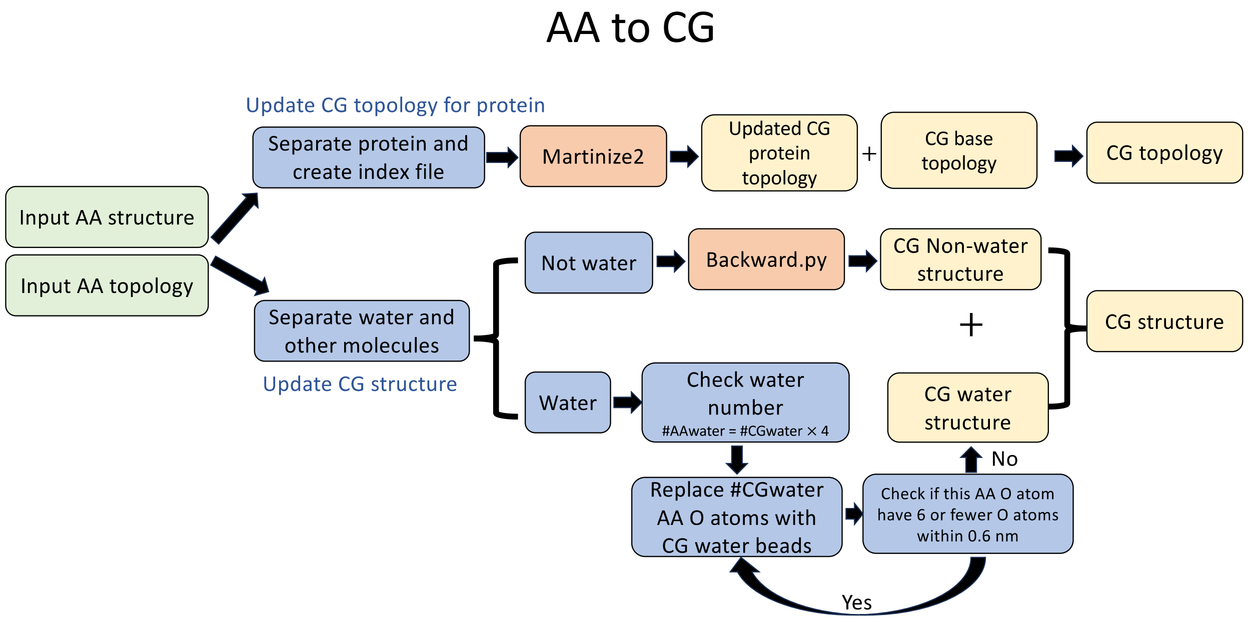


*Figure S1 Workflow of AA-to-CG conversion.*

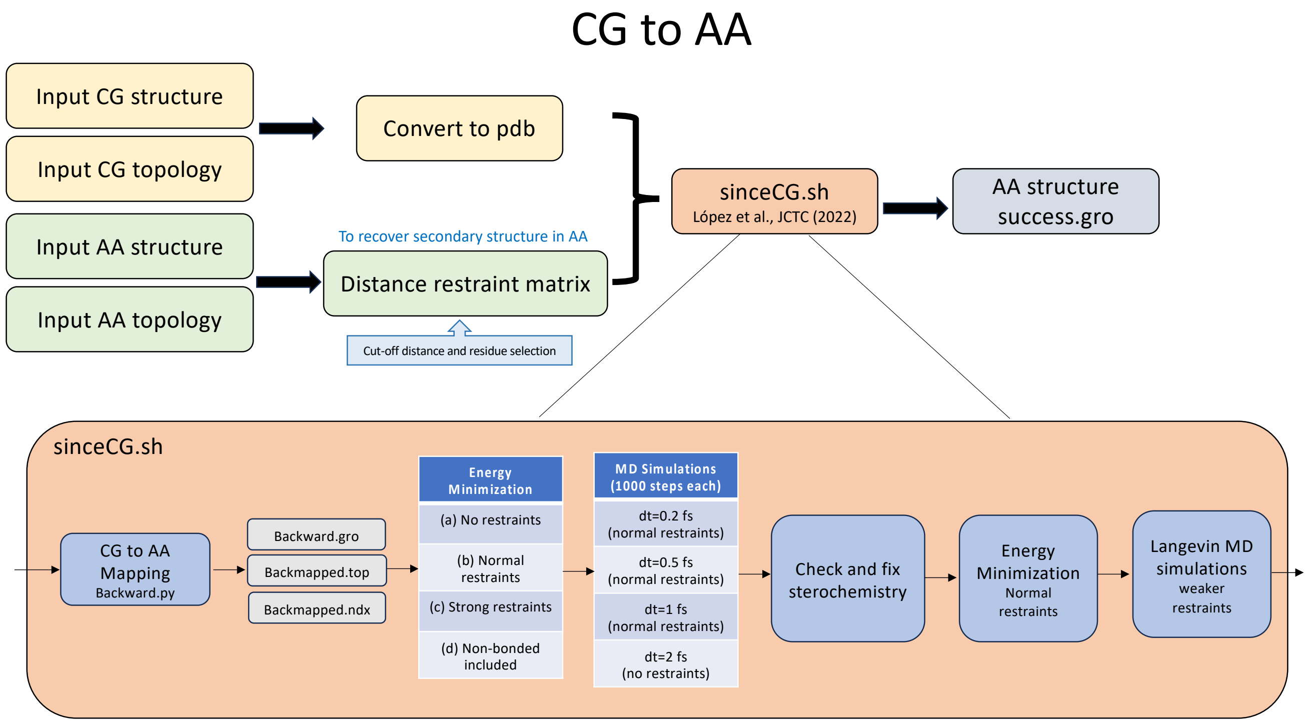


*Figure S2 Workflow of CG-to-AA conversion.*

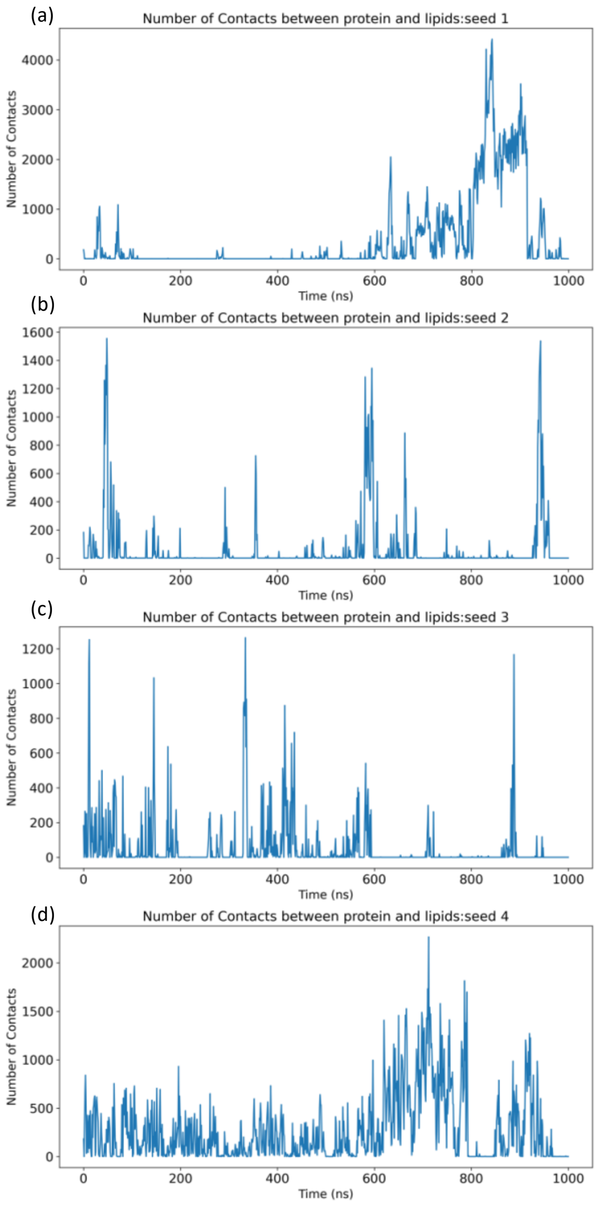


*Figure S3 Time evolution of numbers of contact for atom pairs within 6 Å cutoff radius between protein and lipid membranes in the first AA simulations from 4 independent simulations (a-d) starting with the same starting point as the example run shown in* ***Fig. 3****.*

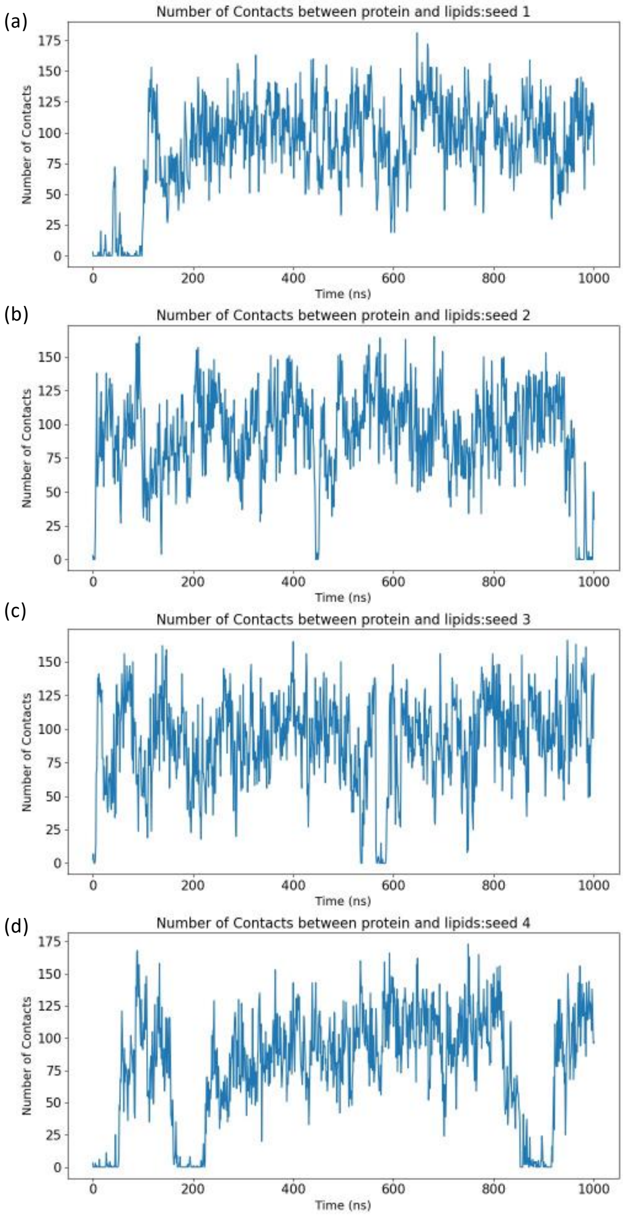


*Figure S4 Time evolution for numbers of contacts within 6 Å cutoff between the fusion peptide and membrane in the CG system respectively from 4 independent simulations (a-d) starting with the same starting point as the example run shown in **Fig. 5**. These independent runs are consistent with the description of the representative run in **Fig.5**, i.e., fusion peptides in the CG system can insert into membranes quickly, but it cannot stay firmly inside the membrane.*

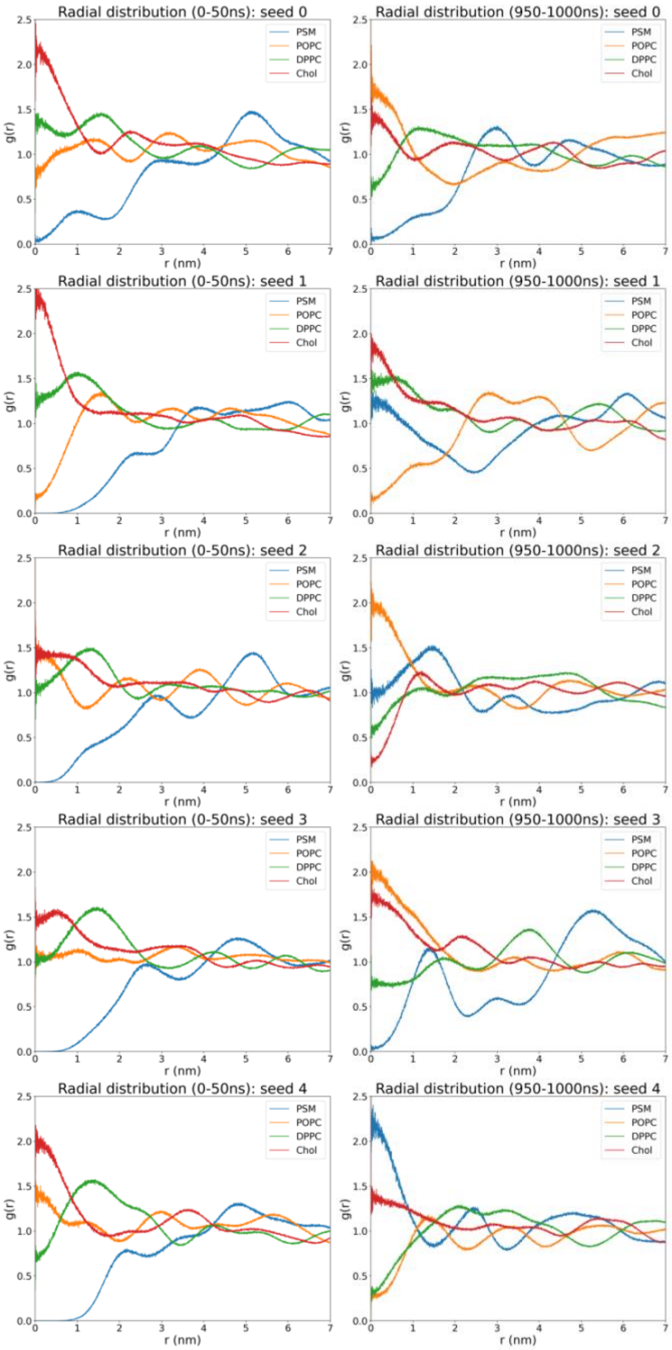


*Figure S5 Radial distribution functions of the fixed helix and lipids at the (left) early stage (0-50 ns) and (right) late stage (950-1000 ns) for the AA1 systems, respectively.*

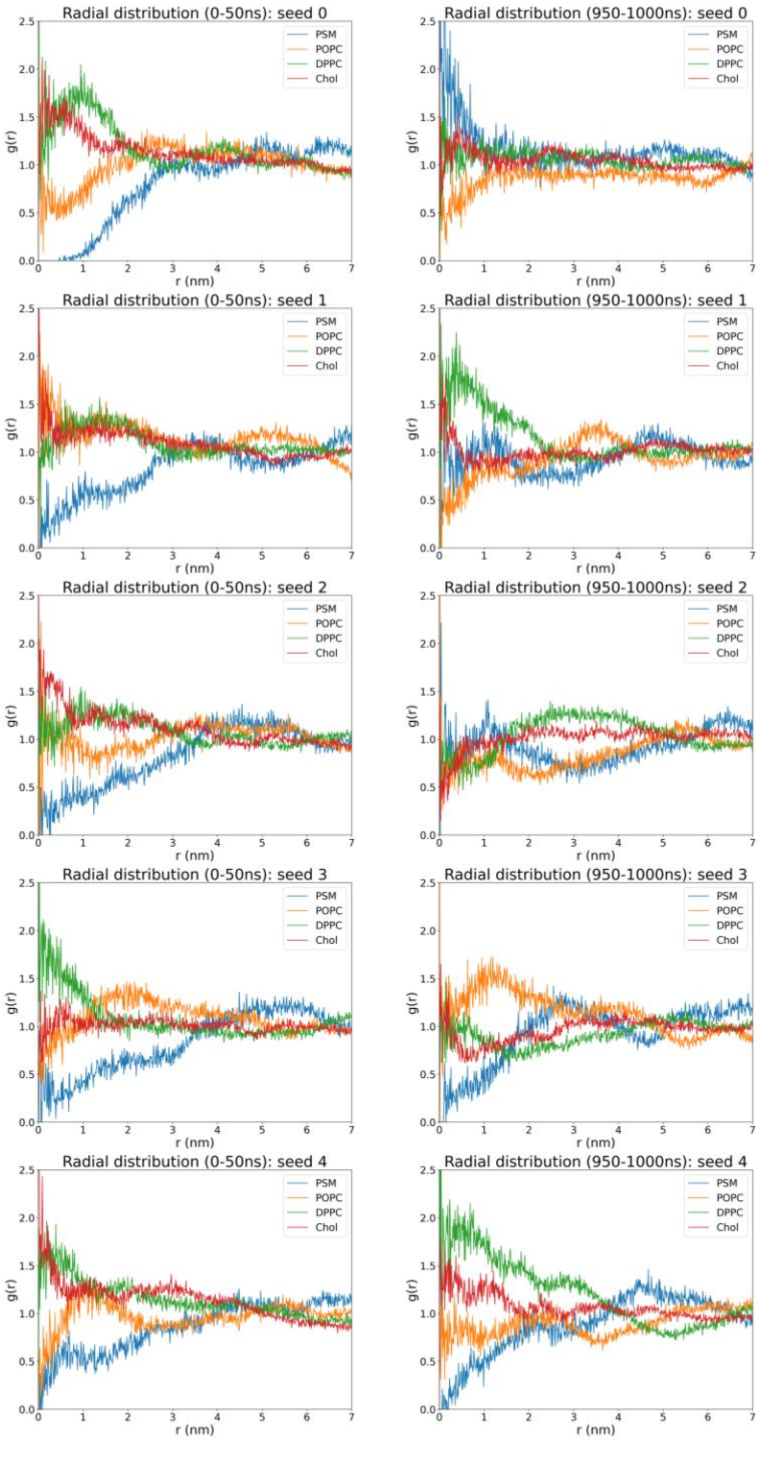


*Figure S6 Radial distribution functions of the fixed helix and lipids at the (left) early stage (0-50 ns) and (right) late stage (950-1000 ns) for the CG systems, respectively.*

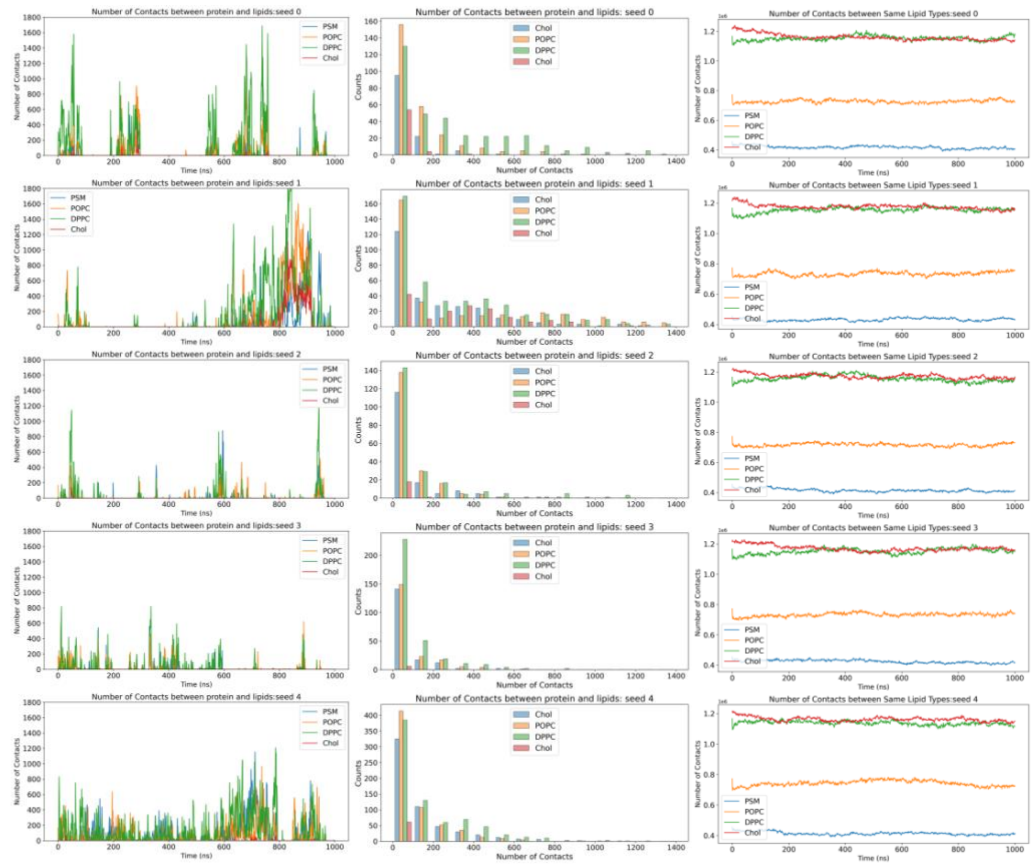


*Figure S7 Time evolution for numbers of contacts within 6 Å cutoff (left) between the fusion peptide and membrane, (middle) histograms for numbers of contacts between protein and each lipid type, and (right) among the same lipid type in the AA1 system respectively from 5 independent simulations.*

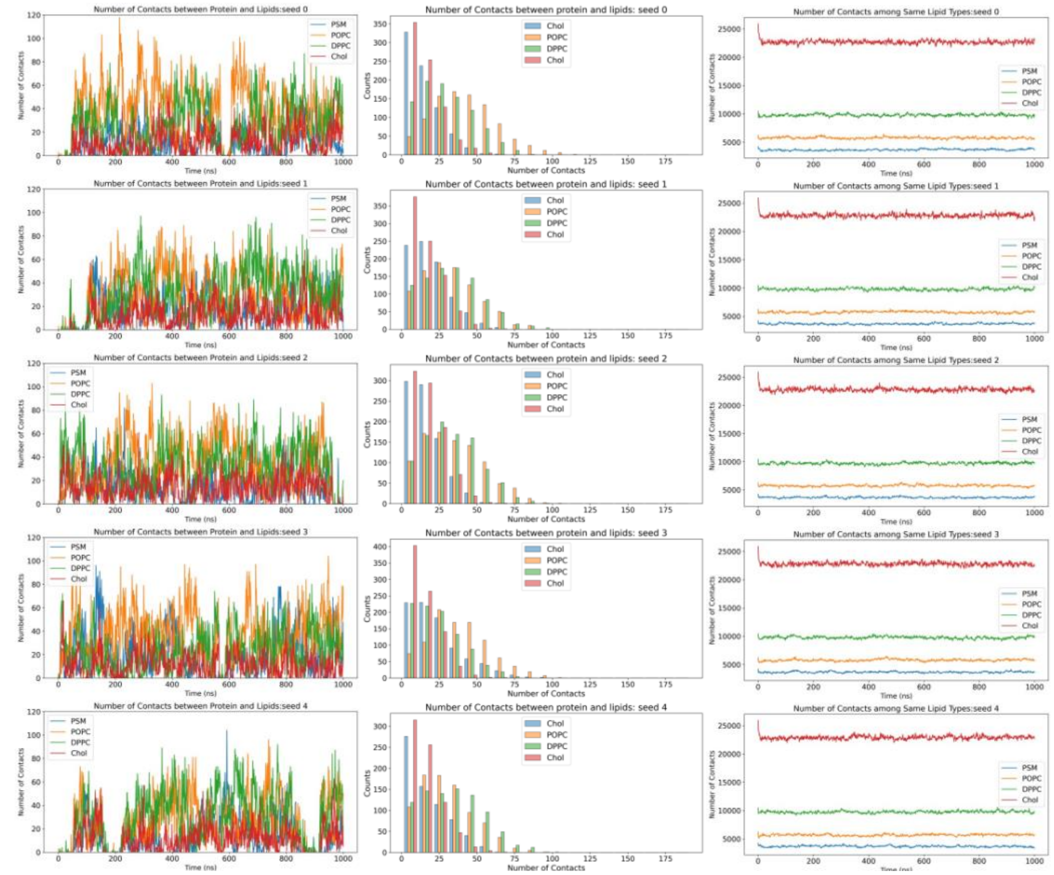


*Figure S8 Time evolution for numbers of contacts within 6 Å cutoff (left) between the fusion peptide and membrane, (middle) histograms for numbers of contacts between protein and each lipid type, and (right) among the same lipid type in the CG system respectively from 4 independent simulations.*

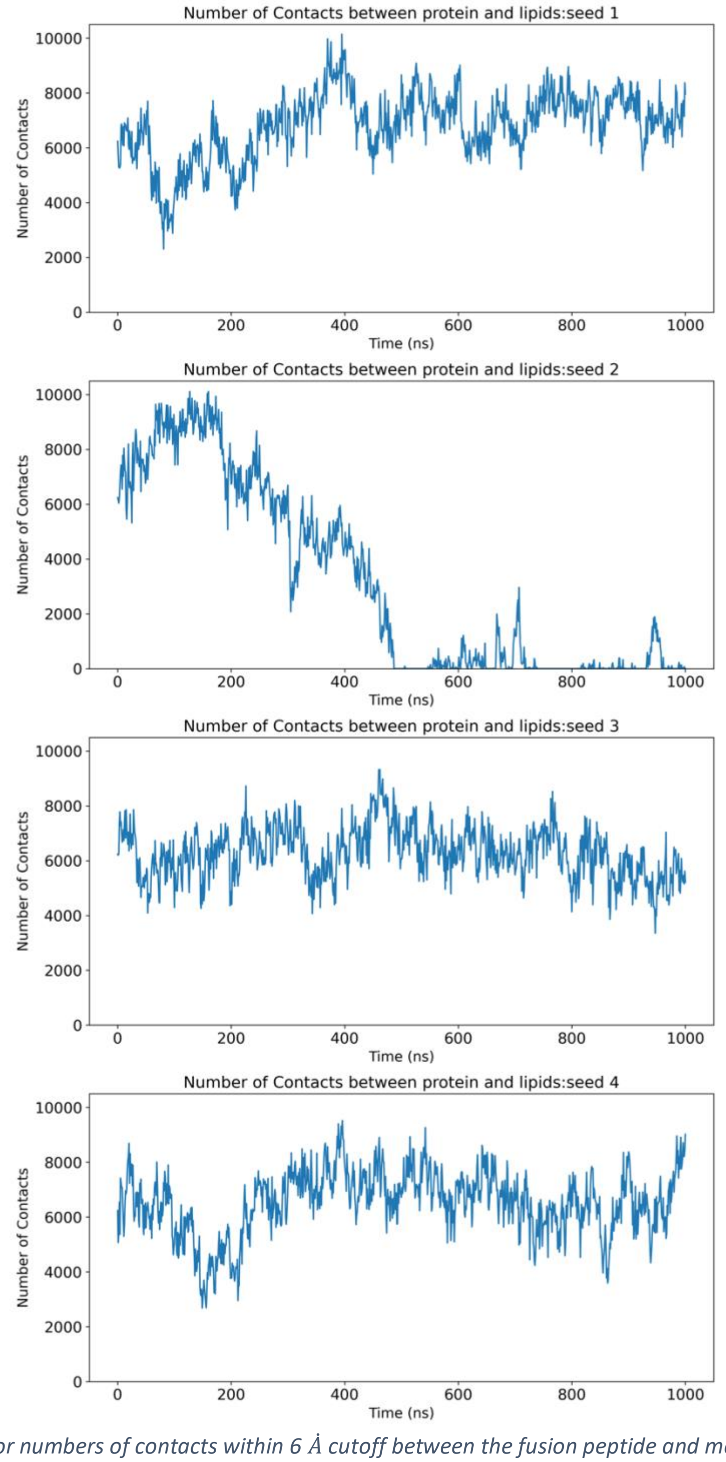


*Figure S9 Time evolution for numbers of contacts within 6 Å cutoff between the fusion peptide and membrane in the AA2 system respectively from 4 independent simulations (a-d) starting with the same starting point as the example run shown in* ***Fig. 8****. These independent runs are consistent with the description of the representative run in* ***Fig.8****.*

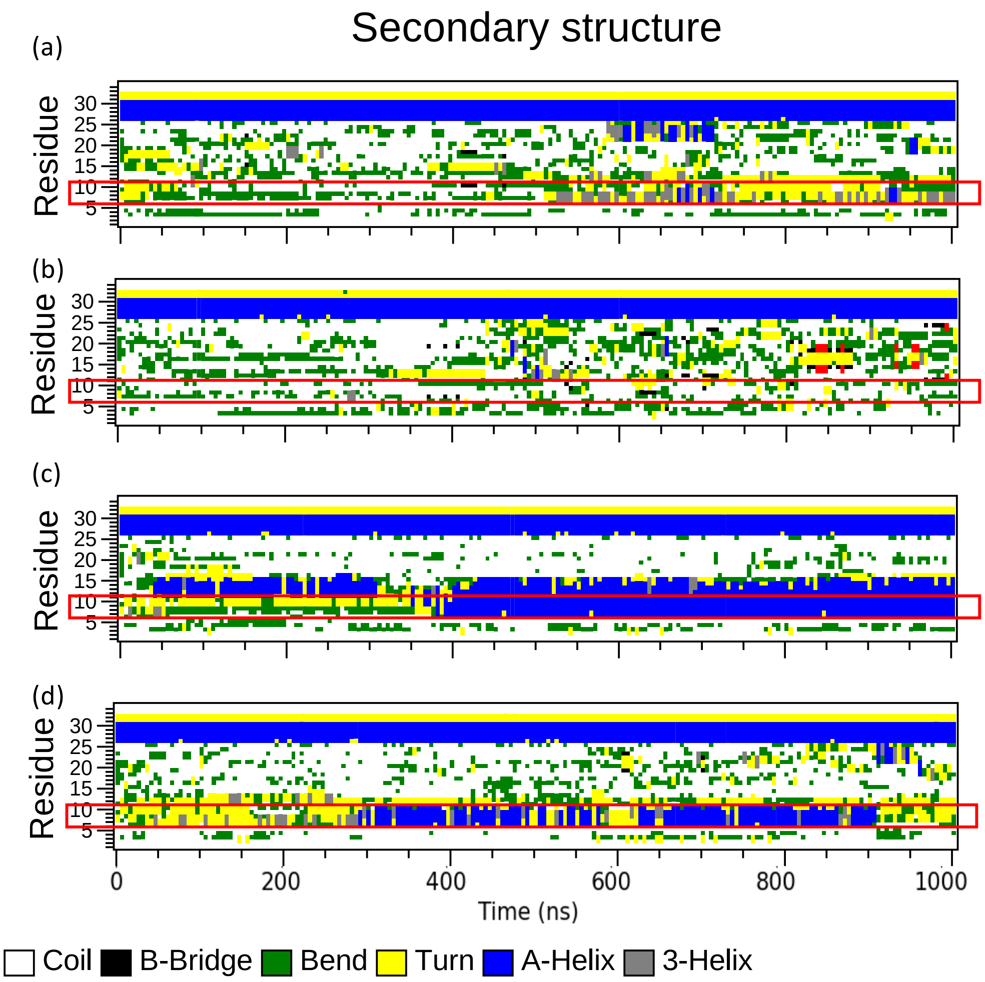


*Figure S10 Time evolution of the secondary structures for each residue in the AA2 run during the 1 μs simulation course from 4 independent simulations (a-d) starting with the same starting point as the example run shown in* ***Figure 8****. In 3 out of these 4 independent runs (a, c, and d), fusion peptides form some types of helices. The only one does not form a helix (b) cannot stay firmly inside the membrane and leave the membrane around 480 ns. The region of initial folding is consistent with the representative run, i.e., near the "AXXXG" motif.*

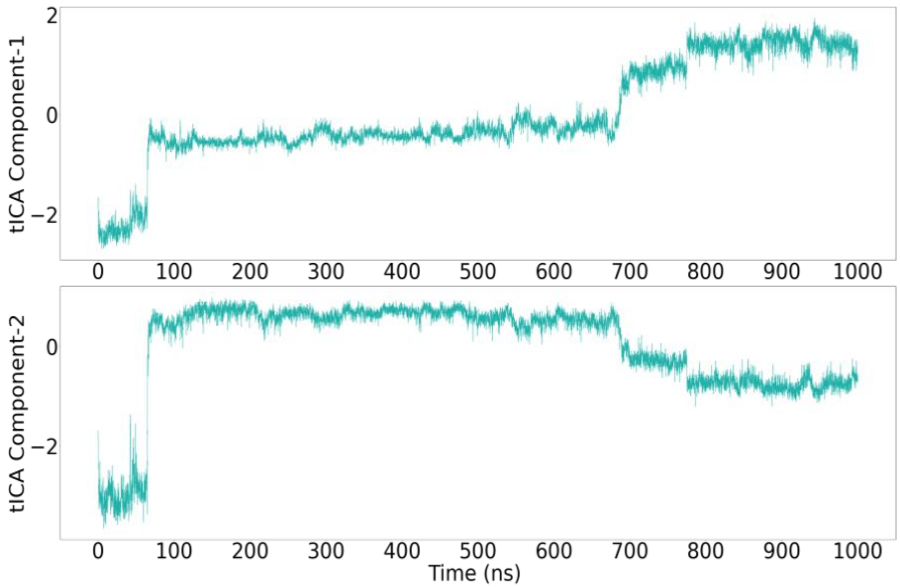

*Figure S11 TICA plots of two components to show the conformational change of the fusion peptide in the AA2 simulation.*

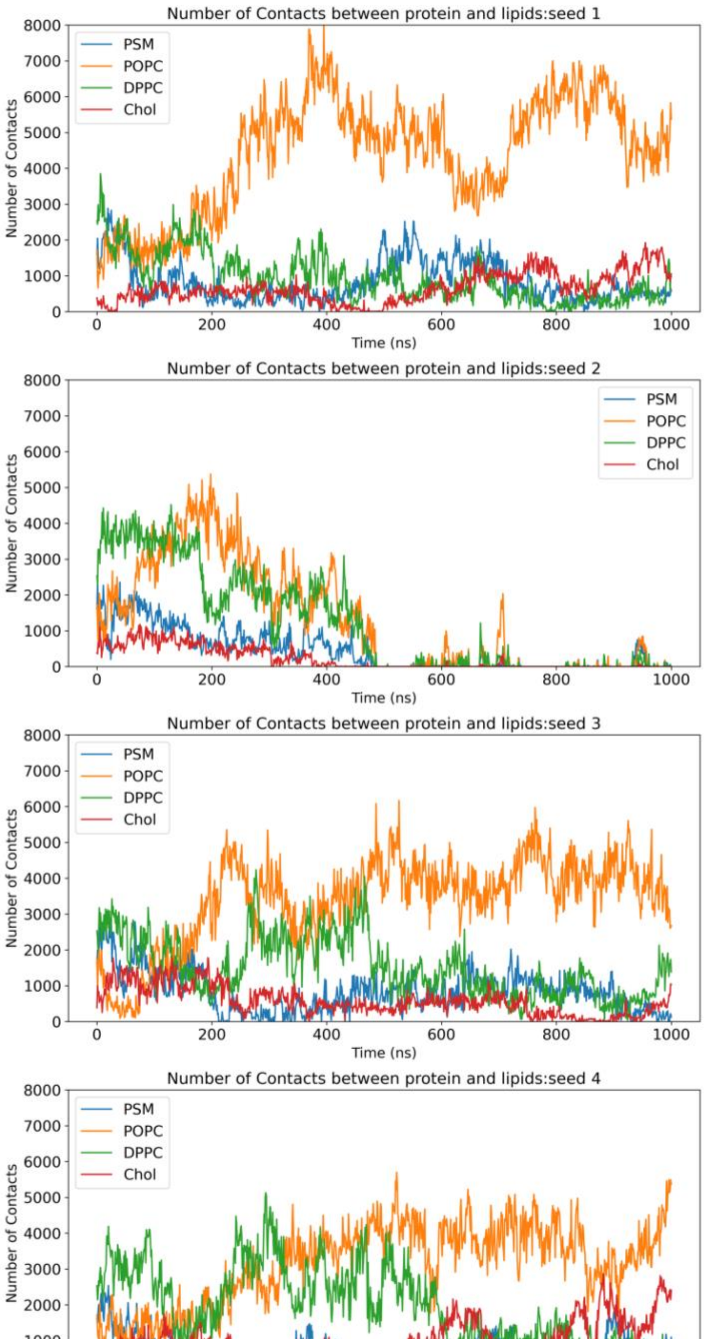


*Figure S12 Time evolution for numbers of contacts within 6 Å cutoff between the fusion peptide and different lipids in the AA2 system from 4 independent simulations.*

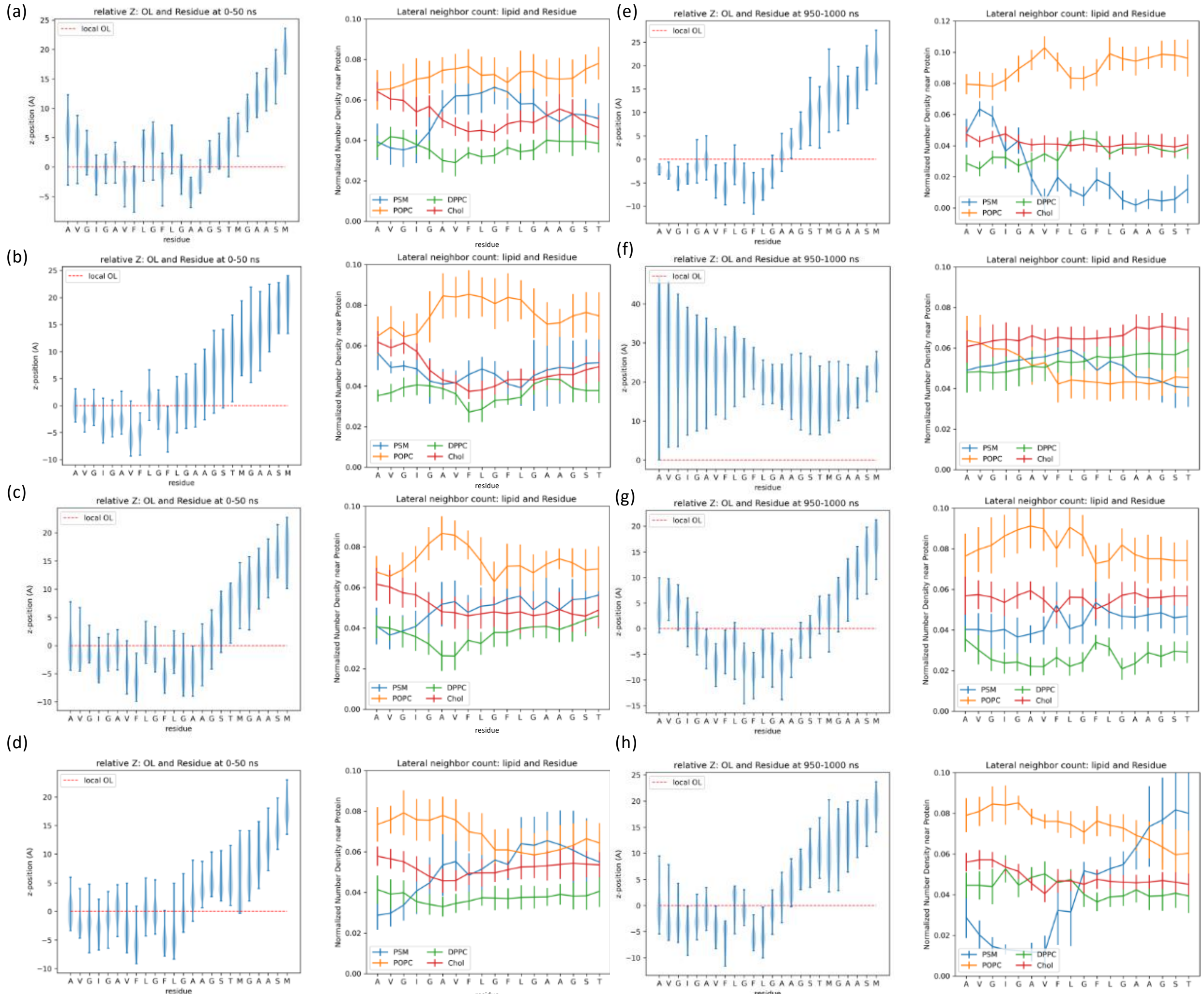


*Figure S13 Average insertion depth of each residue of HIV-1 fusion peptide and normalized lipid distribution averaged from (a-d) the early stage of simulation. (a-d) are calculated from four independent runs starting with the same starting point as the example run shown in* ***Figure 6****. As mentioned before, the fusion peptide in (a) has already left the membrane at this time period. We can observe it from the residue positions in the left panel (b).*

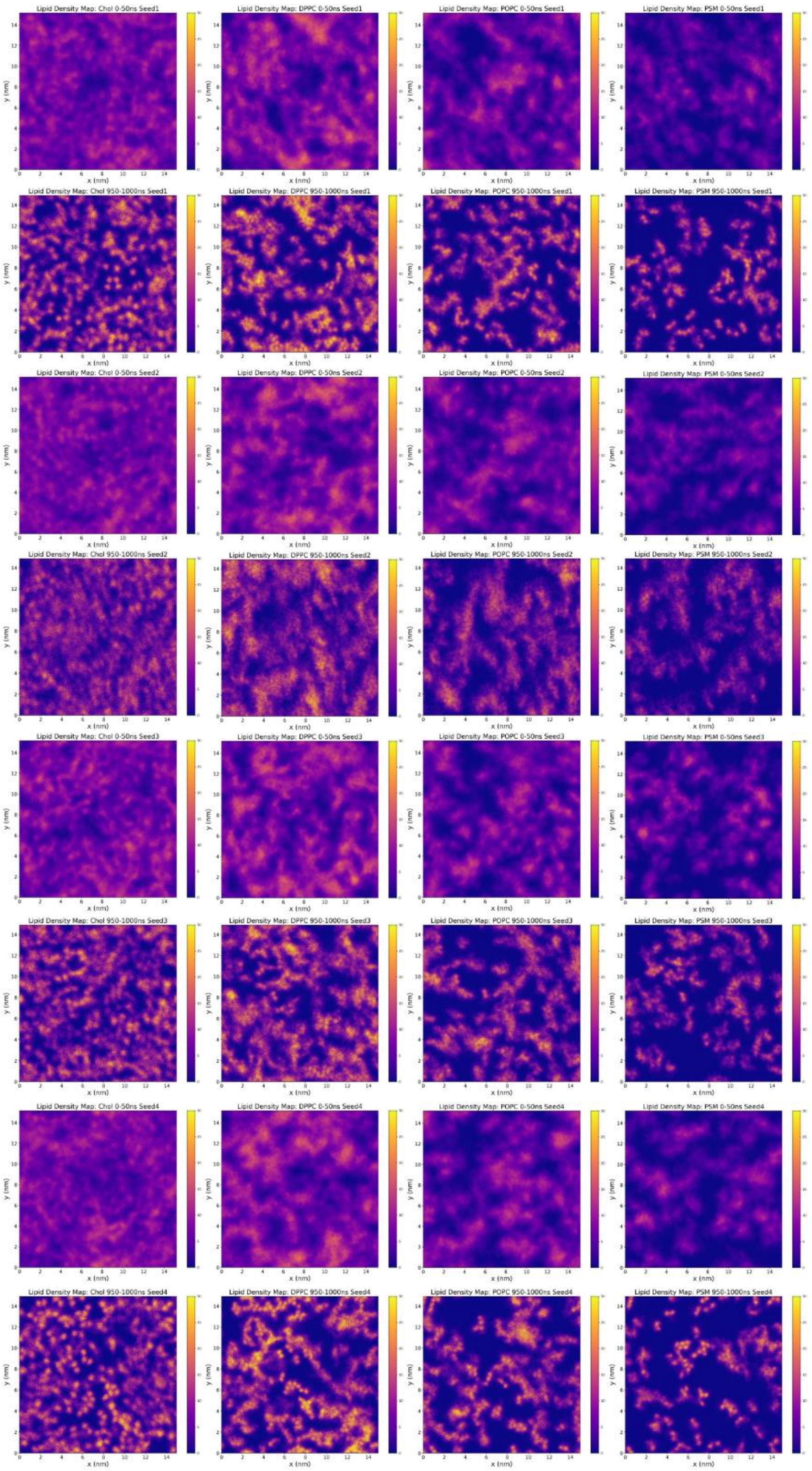


*Figure S14 Average number density maps of lipids on the upper leaflet at the early stage (0-50 ns) and the late stage (950 - 1000 ns), respectively. The membrane-embedded portions (residue 512 to 528) of the fusion peptides are in the center of each density map, but not showing here for simplicity. Other settings are consistent with **Figure 12**.*
